# Chicken Auditory Supporting Cells Express Interferon Response Genes during Regeneration towards Nascent Sensory Hair Cells *In Vivo*

**DOI:** 10.1101/2021.09.21.461299

**Authors:** Amanda Janesick, Mirko Scheibinger, Nesrine Benkafadar, Sakin Kirti, Stefan Heller

**Author notes:** Correspondence (A.J.) or (S.H.).

## Abstract

The avian hearing organ is the basilar papilla that, in sharp contrast to the mammalian cochlea, can regenerate sensory hair cells and thereby recover from complete deafness within weeks. The mechanisms that trigger, sustain, and terminate the regenerative response *in vivo* are largely unknown. Here, we profile the changes in gene expression in the chicken basilar papilla after aminoglycoside antibiotic-induced hair cell loss using RNA-sequencing. The most prominent changes in gene expression were linked to the upregulation of interferon response genes which occurred in supporting cells, confirmed by single-cell RNA-sequencing and *in situ* hybridization. We determined that the JAK/STAT signaling pathway is essential for the interferon gene response in supporting cells, set in motion by hair cell loss. Four days after ototoxic damage, we identified newly regenerated, nascent auditory hair cells that express genes linked to termination of the interferon response. These cells are incipient modified neurons that represent a population of hair cells *en route* towards obtaining their location-specific and fully functional cell identity. The robust, transient expression of immune-related genes in supporting cells suggests a potential functional involvement of JAK/STAT signaling and interferon in sensory hair cell regeneration.

## Introduction

Hearing regeneration in the avian auditory organ, known as the basilar papilla, showcases the power of non-mammalian regenerative capabilities observed in such organisms as chickens, zebrafish and salamanders. Universal regenerative signals shared between these species and their organ systems (e.g., lens, heart, and limb) are appreciated to include nerve dependence, thrombin activation, and immunomodulation (Brockes and Kumar, 2008). Mobilization of immune cells or activation of immune genes is essential for debris removal, extracellular matrix remodeling, and secretion of signals to promote proliferation and wound healing (Julier et al., 2017). The role of immune processes in regulating the pathology of hearing loss in non-regenerating, mammalian systems has also gained traction where inflammation and immune cell infiltration are both seen as protective, but also harmful when linked to fibrosis after sensory hair cell loss (Hu et al., 2018; Rai et al., 2020; Raphael et al., 2007; Wood and Zuo, 2017). While both the avian basilar papilla and the mammalian cochlea invoke an immune response after damage (Hirose et al., 2017; Warchol, 1997), one happens in the context of regeneration and the other does not. In the regenerative avian sensory epithelium, macrophage infiltration and up-regulation of immune genes has been reported (Matsunaga et al., 2020; Warchol, 1997; Warchol et al., 2012).

Previous reports have investigated changes in gene expression following hair cell damage in the chicken basilar papilla and utricle (**Table S1**). These studies employed an *in vitro* culture model and bulk RNA-sequencing methods. *In vitro* models are valuable for immediate and homogeneous application of drugs and compounds but are limited by the lack of surrounding tissue and the tendency of explants to lose complex organ structure. Moreover, bulk RNA-sequencing techniques lack the resolution to distinguish whether up-regulated genes, such as immune genes, emanate from immune cells or otic-derived sensory epithelial cells. We recently unveiled a surgical model for local infusion of the ototoxic drug sisomicin *in vivo* via the posterior semi-circular canal of the chicken ear (Benkafadar et al., 2021; Janesick et al., 2021a -- see Supplemental Files section for preprint). This method yields rapid extrusion of hair cells and temporal synchrony of the regenerative response, which is essential for transcriptomic analyses. Since this surgical method is an *in vivo* model, we are confident that it represents a relatively complete hair cell regeneration model based on 3-week data shown in the present study, as well as historical reports that deafened songbirds can relearn vocal mimicry after damage (Bermingham-McDonogh and Rubel, 2003; Ryals et al., 2013).

Leveraging recent undamaged homeostatic “baseline” single cell data of the avian basilar papilla (Janesick et al., 2021b), we now have a clear roadmap for detecting changes in gene expression occurring after ototoxic insult. We recently utilized the baseline dataset to explore hair cell demise from 12-24 hours after damage (Benkafadar et al., 2021). Our present study aims to investigate the regenerative process at 30, 38, and 96 hours after sisomicin infusion with single cell resolution. We characterize a group of responding supporting cells at 30 and 38 hours and found the most prominent change in gene expression in these cells was linked to the interferon signaling cascade. These responding supporting cells possess gene expression profiles similar to leukocytes, despite being derived from the ectodermal otic placode. We further show that the expression of interferon response genes within the supporting cell layer is linked to JAK activation and that inhibition of JAK/STAT signaling abolishes the upregulation of these genes. At 96 hours, we profile a small group of conspicuous *ATOH1*-expressing, inchoate hair cells, and compare them to mature basilar papilla hair cells as well as nascent vestibular hair cells. We identify distinct markers that are expressed in responding supporting cells as well as in newly regenerated nascent hair cells, including *USP18* which we posit to halt the interferon response, and *CALB2* which was not previously appreciated as an early marker of the regenerative response. Collectively, our results identify immune-like behavior in supporting cells followed by differentiation into nascent, regenerated hair cells.

## Results

### Interferon Response Genes are Expressed in the Chicken Basilar Papilla after Hair Cell Loss In Vivo

We recently developed a surgical method to eliminate all hair cells in the chicken basilar papilla by infusing the ototoxic aminoglycoside sisomicin into the inner ear via canalostomy (Benkafadar et al., 2021; Janesick et al., 2021a). This *in vivo* damage paradigm differs from existing models because it requires only a single application of the ototoxin, and hair cell extrusion happens within 24 hours (Benkafadar et al., 2021; Janesick et al., 2021a), resulting in temporal synchrony of the regenerative response. Forty-eight hours after damage, we dissected the basilar papilla and employed a rapid “cold peeling” method for obtaining pure sensory epithelia, which does not require the application of proteases and time-consuming incubation (Janesick et al., 2021a; Janesick et al., 2021b). This method allows epithelial cells to be harvested and lysed within ten minutes of sacrificing the animal. We infused sisomicin into the left inner ears of seven-day-old chickens (P7) and isolated RNA 48 hours post-surgery (P9) from cold-peeled contralateral control and damaged epithelia. Triplicate samples were submitted for bulk RNA-sequencing (RNA-seq).

RNA-seq reads were annotated using the latest genome information for *Gallus gallus* (GRCg6a), which yielded 96.9% uniquely mapped reads and 22,758 genes. Cells from control and sisomicin-infused basilar papillae clustered into distinct groups and correlated well between the two conditions (**Fig. S1A**). The data were normalized and filtered (Rau et al., 2013; Robinson and Oshlack, 2010) to remove constant and low-abundance transcripts, leaving 11,952 genes (**Fig. S1B-D**). Differential expression analysis with EdgeR (Robinson et al., 2010) revealed 637 genes with log_2_FC ≥ 2 in the sisomicin-infused condition and FDR < 0.05, relative to the control, where FC is fold-change, and FDR is false discovery rate (**Fig. 1A)**. The entire differential expression dataset can be found in **Table S2** and visualized with an interactive Glimma plot (Su et al., 2017) − see Supplemental Files. 995 genes were expressed at log_2_FC < −2 in the sisomicin-infused condition relative to control. Since hair cells are missing in the damaged tissue, we expected that the majority of genes expressed at log_2_FC < −2 would account for missing hair cell genes. We created a heatmap for the top differentially expressed genes of log_2_FC > 5, FDR < 0.05, and log_2_CPM > 5 where CPM is counts per million (**Fig. 1B**). As expected, the lowest expressed genes are hair cell-specific, many of which were recently validated by *in situ* hybridization (Janesick et al., 2021b). This confirms that the sisomicin treatment had eliminated auditory hair cells at 48 hours post-surgery. We previously verified that the number of supporting cells remained unchanged after damage (Benkafadar et al., 2021); however, *TECTA*, a supporting cell gene, was significantly down-regulated. In the absence of hair cells, it is possible that genes required for maintaining the tectorial membrane are deprioritized, perhaps until new hair cells with hair bundles are regenerated.

**Figure 1.**
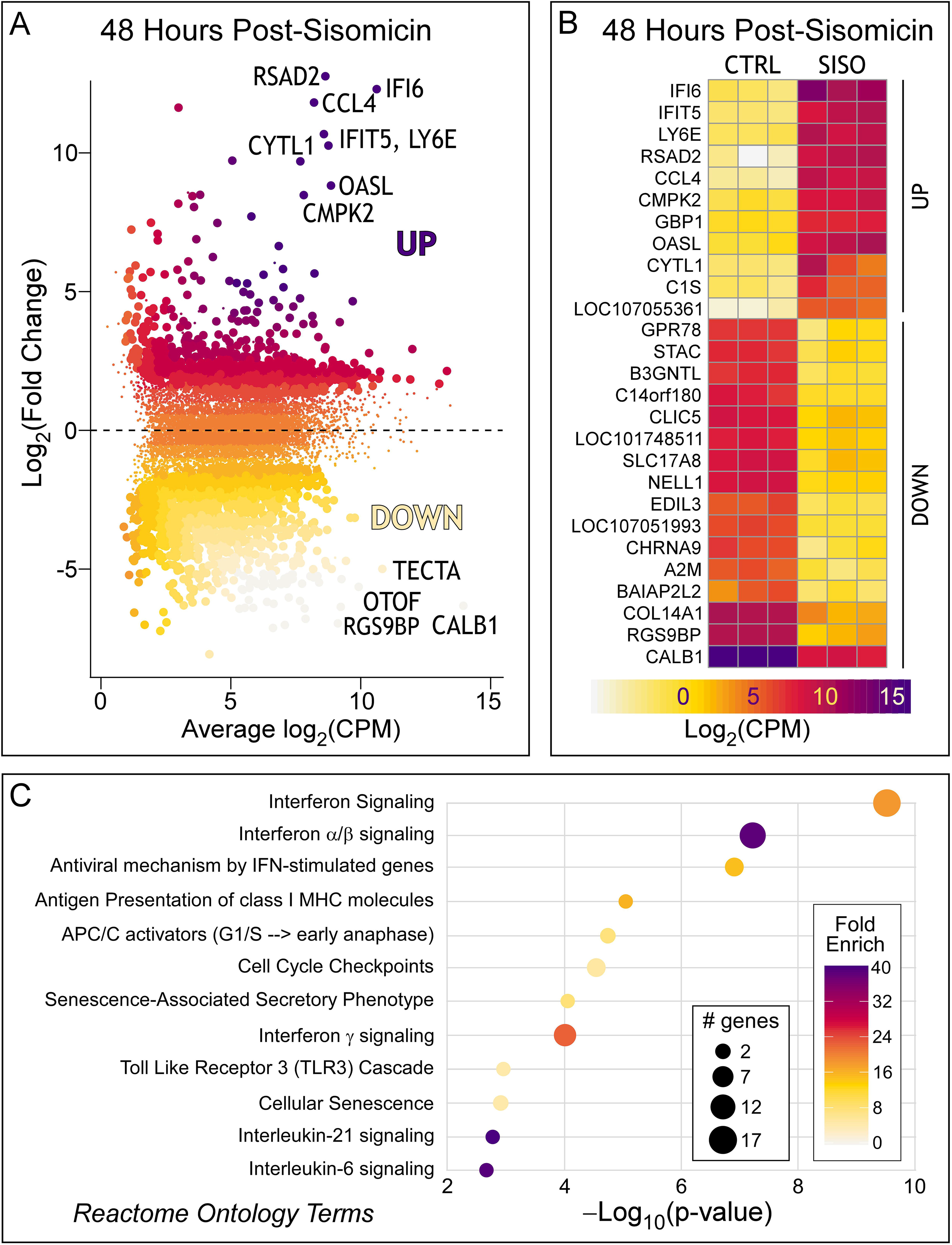
Differentially expressed genes and their ontology at 48 hours post-sisomicin. **(A)** Mean-difference volcano plot highlighting genes (larger dots) with log_2_(fold change) > 2 and FDR < 0.05 (for interactive plot, see *Data Availability* section). Fold change is relative to the contralateral control ear. CPM = counts per million reads. **(B)** Heatmap showing all three biological replicates per treatment, highlighting genes that meet the criteria of log_2_(fold change) > 2, FDR < 0.05, and log_2_CPM > 5. CTRL = Control (undamaged); SISO = Sisomicin damaged. FDR = false discovery rate (Benjamini-Hochberg corrected p-value) **(C)** Gene ontology analysis of sisomicin up-regulated genes using pathfindR analysis tool and Reactome database. See **Table S2** for the full table of genes, their counts, differential expression and associated FDR.

Gene Ontology and Reactome analysis identified a strong correlation between top upregulated genes and an interferon signaling response (**Fig. 1C**). These genes are often activated via the JAK/STAT signaling pathway and are typically associated with macrophages and neutrophils (Goossens et al., 2013; Santhakumar et al., 2017). There is evidence that macrophages infiltrate the chicken basilar papilla after ototoxic insult (Warchol et al., 2012). However, chemical- or genetically-mediated macrophage depletion did not inhibit regeneration of hair cells in the chicken basilar papilla (Warchol et al., 2012) or the zebrafish lateral line (Warchol et al., 2021). A limitation of the bulk RNA-seq approach is that it cannot rule out the possibility that there is infiltration of blood and immune cells into the basilar papilla after sisomicin infusion, which could explain the up-regulation of interferon response genes. We hypothesized that single-cell RNA-seq would provide the cellular resolution for a differential gene expression analysis, allowing us to determine which specific cell type(s) expressed the interferon responding genes after sisomicin-induced hair cell loss.

### Single-Cell RNA-Sequencing of the Regenerating Chicken Basilar Papilla

Pure basilar papilla sensory epithelia were dissociated into individual cells and processed for single-cell RNA-seq as previously described (Ellwanger et al., 2018; Janesick et al., 2021b). In total, we analyzed 1,891 cells at five different time points, each representing three independent clutches of chickens (**Fig. S2**). We chose the 30- and 38-hour timepoints for analysis because this is the peak time of supporting cell S-phase entry post-sisomicin infusion, and hence is affiliated with proliferation of supporting cells (Janesick et al., 2021a). Ninety-six hours post-sisomicin, supporting cells no longer enter S-phase, and the first regenerated MYO7A-positive new hair cells are detectable with immunohistochemistry (Janesick et al., 2021a). Our single-cell sequencing libraries captured 17,557 expressed genes. Data quality control flagged 3,987 low-abundance genes and 670 information-poor cells which were removed from downstream analyses. The remaining 1,221 cells were normalized with SCnorm (Bacher et al., 2017) and log_2_-transformed (see *Data Availability* section for normalized count matrix).

Dimensionality reduction using CellTrails (Ellwanger et al., 2018) revealed seven major epithelial cell groups (**Fig. 2A**) and 11 clusters (**Fig. 2B**). Hair cells were identified by the marker *TMEM255B* (**Fig. 2C**; Janesick et al., 2021b). At the 30- and 38-hour timepoints, we observed compromised hair cells that expressed *TRIM35* (**Fig. 2D**, Benkafadar et al., 2021). These hair cells were also *TMC2*-high and *TMC1*-low (**Fig. S3**), suggesting that they were derived from the distal end of the basilar papilla, where the least sensitive hair cells to sisomicin damage are located (Janesick et al., 2021a). *TRIM35*-positive distal hair cells are likely to mount a repair response and represent a group of hair cells that survived the sisomicin treatment. At 96 hours, *ATOH1*-positive hair cells segregated distinctly (**Fig. 2E**) – we classified these as newly regenerated hair cells. Subclusters of tall and short hair cells were identified by marker genes *CXCL14* and *C14orf180* (**Fig. 2F**), respectively, explored in detail in Janesick et al., 2021b. Homogene and supporting cells were distinct populations marked by *LRP2* and *GSTT1L*, respectively (**Fig. 2G, H**; Janesick et al., 2021b). 30- and 38-hour post-sisomicin cells harbored distinct groups of responding supporting cells and homogene cells (**Fig. 2I**). Leveraging knowledge from the bulk RNA-seq data (**Fig. 1, Table S2**), the responding supporting and homogene cells expressed *IFI6*, an interferon-response gene (**Fig. 2I**). Subclusters of medial and lateral supporting cells were identified by marker genes *LCAT* and *NTN4L* (**Fig. 2J**; Janesick et al., 2021b). Red and white blood cells were identified by *HBA1* and *PTPRC*, respectively (**Fig. 2K, L**). Male- and female-derived cells were present in approximately equal numbers (♀= 586; ♂ = 635; **Fig. 2M**), as indicated by the ubiquitous female transcript derived from the *HINTW* gene (Nagai et al., 2014). Annotation for the cell types within each timepoint was visualized with pie charts in **Fig. S2** and quantified in **Table S3**.

**Figure 2.**
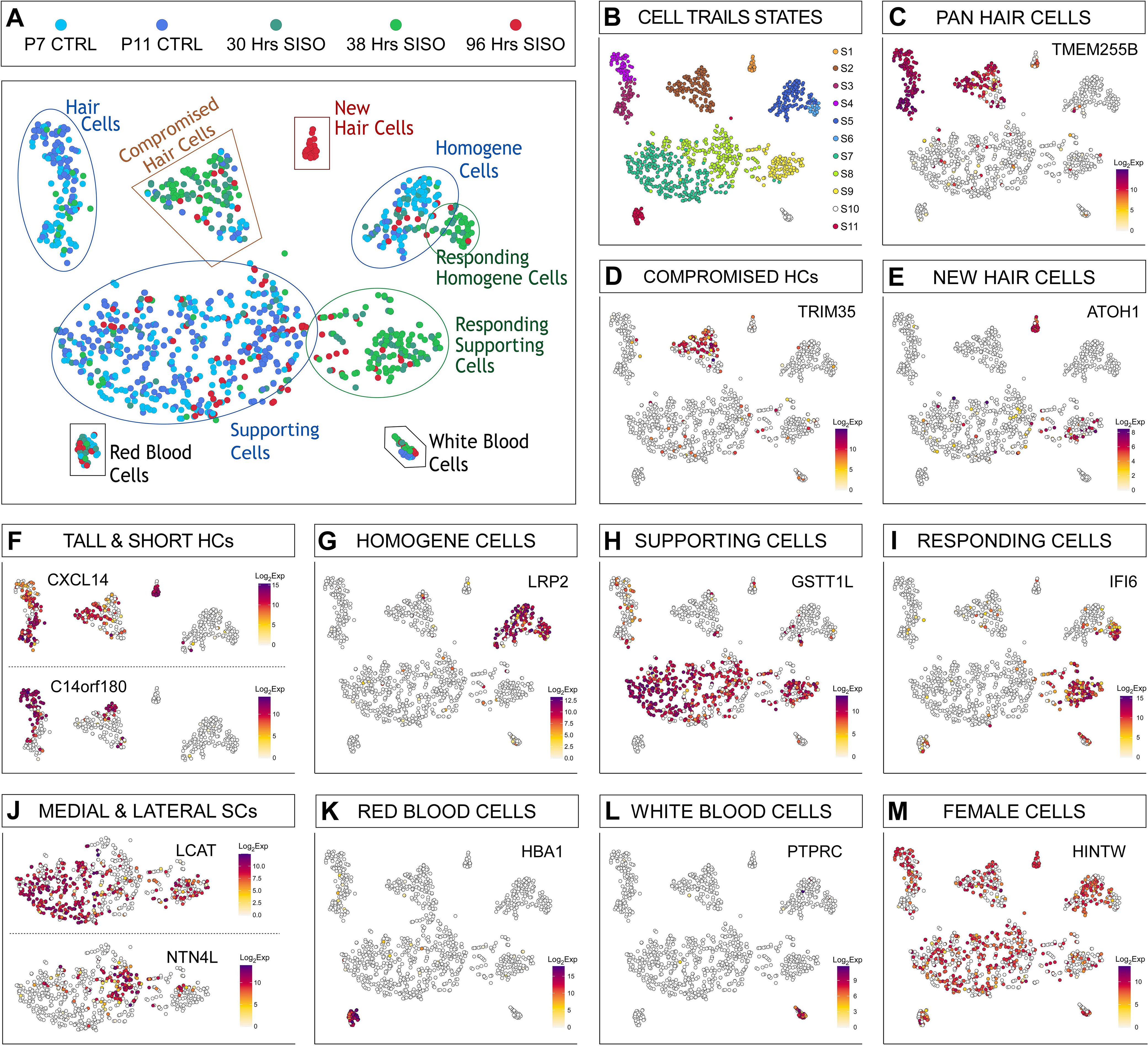
Single cell RNA-Sequencing of the regenerating chicken basilar papilla. T-distributed stochastic neighbor embedding (tSNE) plots of all profiled cells representing 5 timepoints, clustered with CellTrails. **(A)** Seven major epithelial cell groups are outlined. Control (undamaged); SISO = Sisomicin damaged. **(B)** CellTrails clustering reveals eleven states. **(C, F-H, J)** All baseline supporting cell, hair cell, and homogene groups were identified by markers from Janesick et al., 2021b. HCs = hair cells; SCs = supporting cells. **(D)** Compromised hair cells marked by *TRIM35* were defined in Benkafadar et al., 2021. **(E)** New hair cells express *ATOH1*. **(F)***CXCL14* is a tall hair cell marker and *C14orf180* is a short hair cell marker. **(I)** Responding supporting cells express *IFI6*, which was identified in the bulk RNA-seq analysis (see **Fig. 1** and **Table S2**). **(J)** *LCAT* is a medial (a.k.a. neural) and *NTN4L* is a lateral (a.k.a. abneural) supporting cell marker. **(K, L)** Hematocytes. **(M)** *HINTW* tracks female cells. Scalebar = log_2_ transformed, normalized expression counts.

### Interferon Response Genes are Robustly Upregulated in Supporting Cells after Hair Cell Damage

We hypothesized that single-cell RNA-seq would provide the resolution to distinguish whether interferon-inducible genes are expressed by infiltrating immune cells or by sensory epithelial cells. The single cell analysis showed that the major responding cell group, signified by *IFI6* (**Fig. 2I**), also expressed the supporting cell marker *GSTT1L* (**Fig. 2H**). Thus, we inferred that supporting cells were the primary cell type showing upregulation of interferon response genes after damage. A small group of homogene cells, identified with the marker *LRP2*, also expressed *IFI6* (**Fig. 2G, I**). We conducted differential gene expression analysis on the responding supporting cell cluster (State S9 in **Fig. 2B**), comparing it to all other supporting cells (States S7∪S8 in **Fig. 2B**). This analysis revealed 43 up-regulated genes (**Fig. 3A** and **Table S4**) using an FDR cut-off of 0.01 and log_2_FC > 2. We found minimal down-regulation of most supporting cell genes in the responding group, except for a modest reduction in *TECTA*, *SERPINF2*, *HEY2*, *FGFR3*, and *TIMP3* (**Table S4**). Gene ontology analysis corroborated the bulk RNA-seq results, with the top up-regulated terms falling into the category of interferon signaling response and modulation of an immune response (**Fig. S4A**).

**Figure 3.**
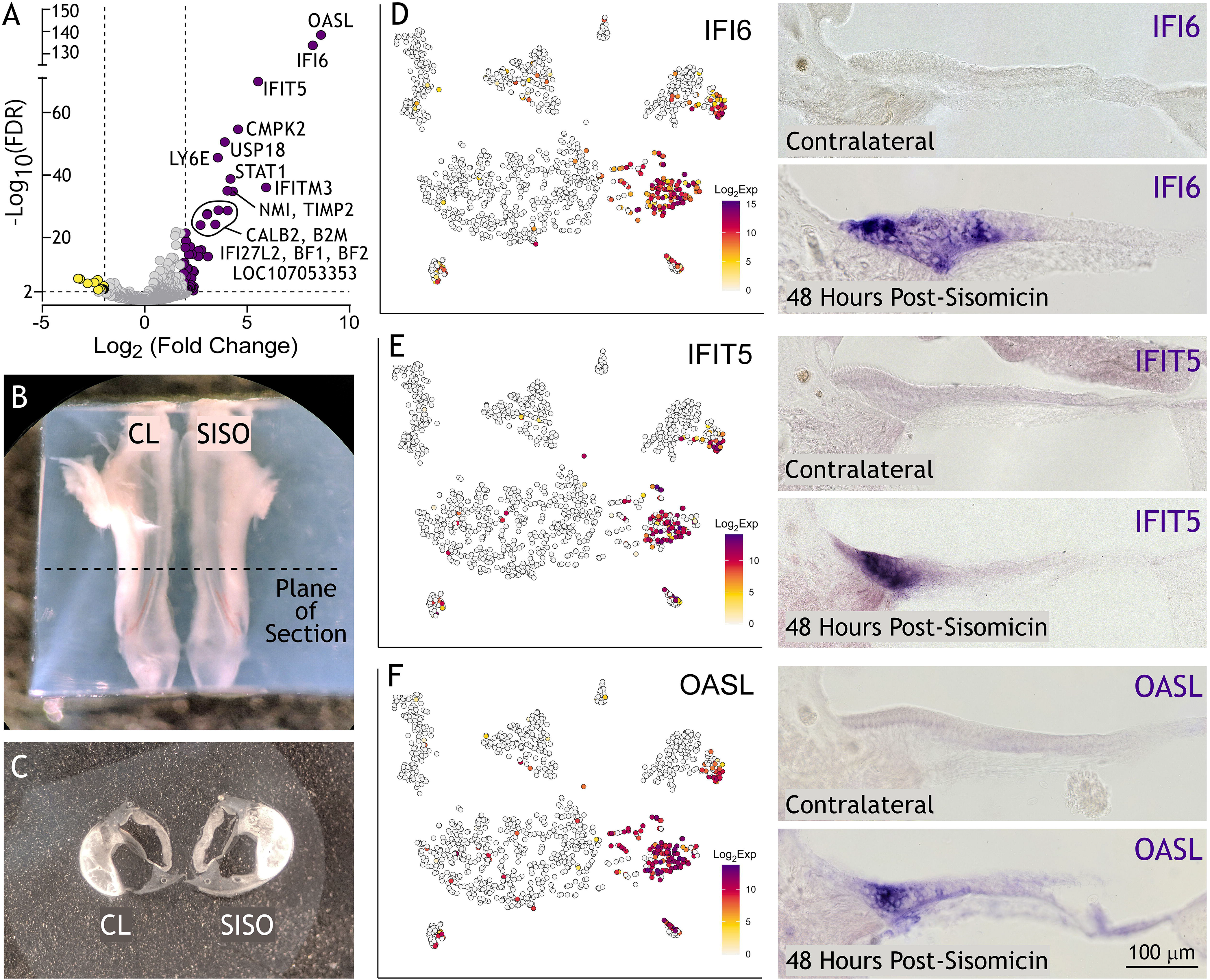
Interferon response genes are highly up-regulated in the responding supporting cell cluster. **(A)** Volcano plot illustrating genes expressed in responding supporting cells (purple dots) at least 4-fold higher and significantly different (FDR < 0.01) compared with genes expressed in supporting cells (yellow dots). **(B)** Agarose embedded basilar papilla from contralateral (CL) and sisomicin-infused (SISO) ears. The dotted line indicates the tonotopic middle of the basilar papilla from where vibratome sections are taken. **(C)** Resulting vibratome sections of CL and SISO basilar papillae. Ejected hair cells are visible in the SISO section. **(D-F)** tSNE plots (left panels) project log_2_ transformed, normalized expression counts for IFI6, IFIT5, and OASL. In situ hybridization (right panels) showing corresponding mRNA expression in P9 transverse sections, 48 hours post-sisomicin damage. Hair cells are noticeably gone in the sisomicin-infused basilar papillae sections.

Next, we validated the interferon response genes using colorimetric *in situ* hybridization. Since this method is not inherently quantitative, we took appropriate measures for the side-by-side comparison of contralateral control versus the sisomicin-infused ear. We embedded the contralateral and sisomicin basilar papillae next to each other in agarose to ensure that transverse vibratome sections would come from the same tonotopic region (**Fig. 3B, C**) and processed both sections together for *in situ* hybridization. *IFI6*, *IFIT5*, and *OASL* were expressed robustly within the supporting cell layer of basilar papillae from sisomicin-infused ears (**Fig. 3D-F**). To test whether immune cells contributed to the interferon response, we immunolabeled basilar papilla sections for the macrophage marker, TAP1, at 48 hours post-sisomicin infusion (**Fig. S4B**). The supporting cell layer was actively proliferating at this time, as indicated by the incorporation of 5-ethynyl-2′-deoxyuridine (EdU), but was devoid of TAP1-positive cells (**Fig. S4B**). We concluded that infiltration of macrophages into the basilar papilla sensory epithelium was undetectable after sisomicin-induced hair cell loss. We noted a subcluster of responding homogene cells, entirely made up of cells from the 30- and 38-hour time points (**Fig. 2A, G; Fig. 2B** – state S6**; Table S3; Fig. S2**). Differential expression of this responding group (State S6 in **Fig. 2B**) versus control homogene cells (State S5 in **Fig. 2B**) revealed a similar collection of genes that we identified in responding supporting cells, belonging to the interferon response pathway (**Fig. S4C, Table S5**). *In situ* hybridization revealed strong expression of *IFI6* mRNA in homogene cells at 48 hours post-sisomicin damage (**Fig. S4D**).

### Responding Supporting Cells Use Distinct Regenerative Mechanisms

Medial and lateral supporting cells employ different mechanisms to restore lost hair cells (Cafaro et al., 2007). We evaluated our damage/regeneration model at three weeks post-sisomicin with continual exposure to EdU to detect cells undergoing S-phase during the proliferative window of 30-80 hours (Janesick et al., 2021a). Virtually all regenerated hair cells on the medial/neural side of the epithelium displayed EdU-positive nuclei, leading us to infer that they emerged nearly exclusively from mitotic events (**Fig. 4A, A’, B**). In contrast, the lateral/abneural side of the epithelium was regenerated mostly via phenotypic conversion, where a supporting cell morphs into a hair cell without dividing first (**Fig. 4A, A’, B**). These results are comparable with previous observations conducted at six days post-damage, calculating that 81% of medial (tall) and 34% of lateral (short) hair cells arise from mitotic division (Cafaro et al., 2007). We quantified ten basilar papilla sections at three weeks post-sisomicin (**Fig. 4C, D**) using three-dimensional rendering in syGlass (Pidhorskyi et al., 2018) and found that 87% of tall and 21% short hair cells incorporated EdU (**Fig. 4D**). 59% of medial and 4% of lateral supporting cells incorporated EdU (**Fig. 4D**). We observed higher hair cell density on the medial/neural side of the epithelium at three weeks, compared to the more sparsely populated regenerated short hair cells, consistent with prioritization of mitotic tall hair cell restoration.

**Figure 4.**
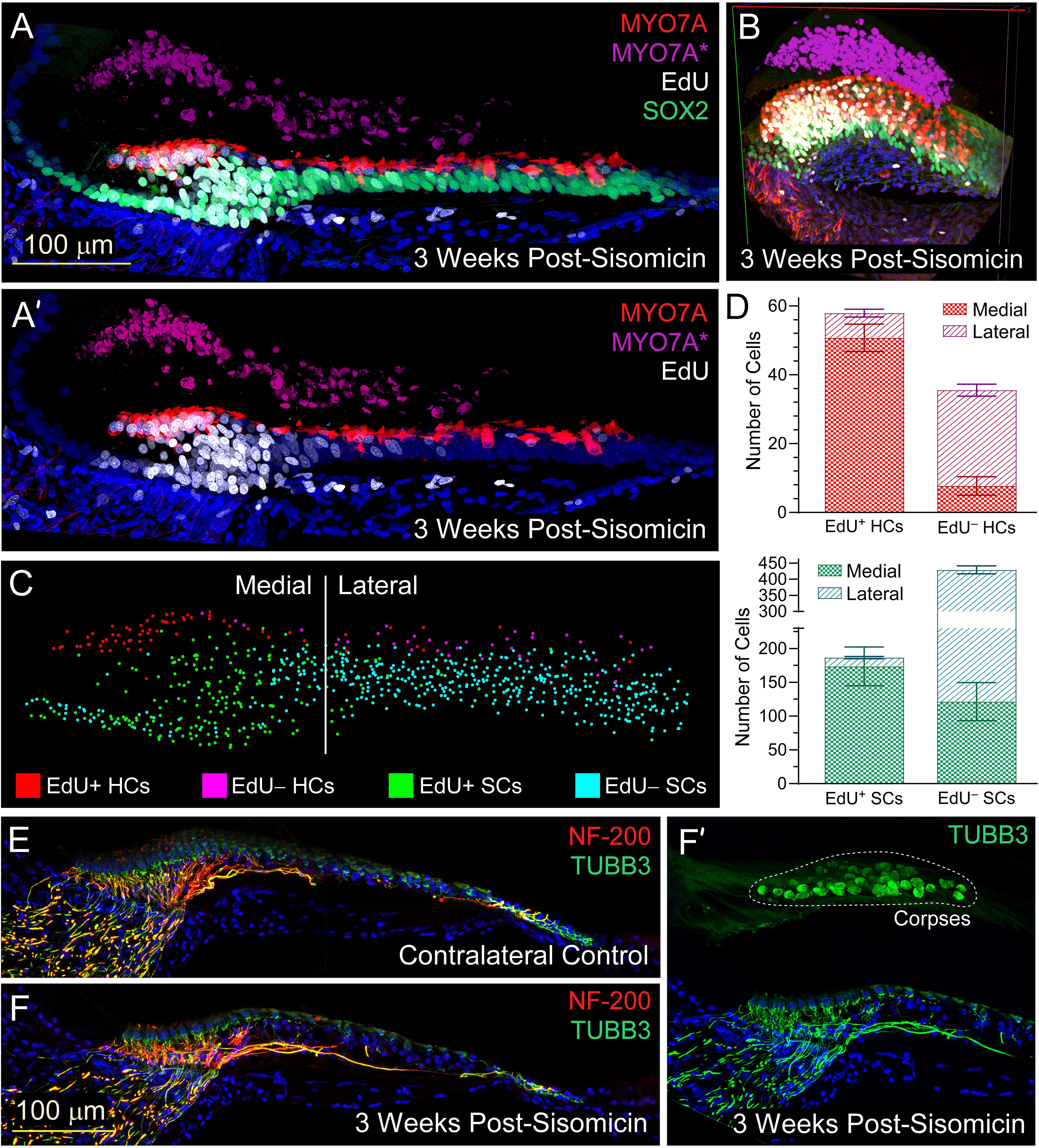
Differential regenerative strategies revealed and quantitated at 3 weeks post-sisomicin. Chickens were injected with EdU for two consecutive days, every 6 hours, during the proliferative window, then sacrificed at 3 weeks. **(A-B)** Immunohistochemistry for MYO7A in red (new hair cells), SOX2 in green (supporting cells), EdU in white (cells which have proliferated), and DAPI in blue (cell nuclei) at three weeks post-sisomicin. MYO7A-positive hair cell corpses are false colored in purple to distinguish them from newly regenerating hair cells in red. **(A’)** SOX2 green channel is removed to clearly visualize EdU staining. **(B)** 3D Visualization-assisted analysis (Vaa3D) image of 3 weeks post-sisomicin basilar papilla. **(C)** The section pictured in (A) was manually quantitated in 3D virtual reality using syGlass software. **(D)** Cell counts from a total of 10 sections were quantitated as shown in (C) and tabulated in bar graphs. Error bars show the standard deviation across the 10 sections. **(E, F, F’)** Immunohistochemistry for Neurofilament-200 (NF-200) in red and β-Tubulin III (TUBB3) in green, which both mark sensory ganglia, and DAPI in blue (cell nuclei). **(E)** Contralateral control. **(F)** Three weeks post-sisomicin. **(F’)** Same image as F but zoomed out to see hair cell corpses in the tectorial membrane, which confirms that these are damaged specimens.

We attempted to subcluster the responding group of supporting cells (State S9 in **Fig. 2B**) further but could not distinguish medial and lateral cells based on markers identified in Janesick et al., 2021b (**Fig. 2J**). It should be noted that such clustering, even of control cells, is not robust because supporting cells are relatively homogeneous, in contrast to the distinct major hair cell subtypes, the tall and short hair cells. Intriguingly, the *in situ* hybridizations in **Fig. 3D-F** revealed higher expression of interferon response genes on the medial side of the epithelium, coincident with EdU staining (**Fig. 4A, B; Fig. S4B**; Janesick et al., 2021a).

### The JAK/STAT Pathway is Essential for Upregulation of Interferon Response Genes in Supporting Cells

We next asked whether JAK/STAT signaling was required to upregulate the genes identified as belonging to the interferon pathway. Ruxolitinib is a dual JAK1 and JAK2 small-molecule inhibitor (Quintás-Cardama et al., 2010). We cultured peeled sensory epithelium in the presence or absence of sisomicin +/− ruxolitinib (RUX) overnight. Then sisomicin was washed away with Medium 199, RUX was refreshed, and the epithelium incubated for six hours with RUX only. mRNA was extracted and processed for qPCR to assess expression of selected interferon-response and control genes. Sisomicin (+/− RUX) reduced the expression of hair cell markers *SLC34A2* and *TMEM255B*, which was expected since hair cells die in the presence of ototoxins (**Fig. 5A**). Expression of supporting cell markers *ZBTB20*, *TECTB*, *TMSB4X*, and *OTOGL* remained constant and unresponsive to chemical treatment (**Fig. 5B**). Sisomicin-induced hair cell loss caused strong upregulation of mRNAs encoding *IFI6*, *IFIT5*, *OASL, USP18, CCL4, RSAD2*, and *LY6E* (**Fig. 5C**). RUX blunted this response by 80-99 percent across all genes and three biological replicates with the exception of CCL4, which was unaffected by RUX (**Fig. 5C**). RUX alone generally did not diminish interferon genes compared to controls. We concluded that the majority of interferon response genes induced after damage require JAK signaling.

**Figure 5.**
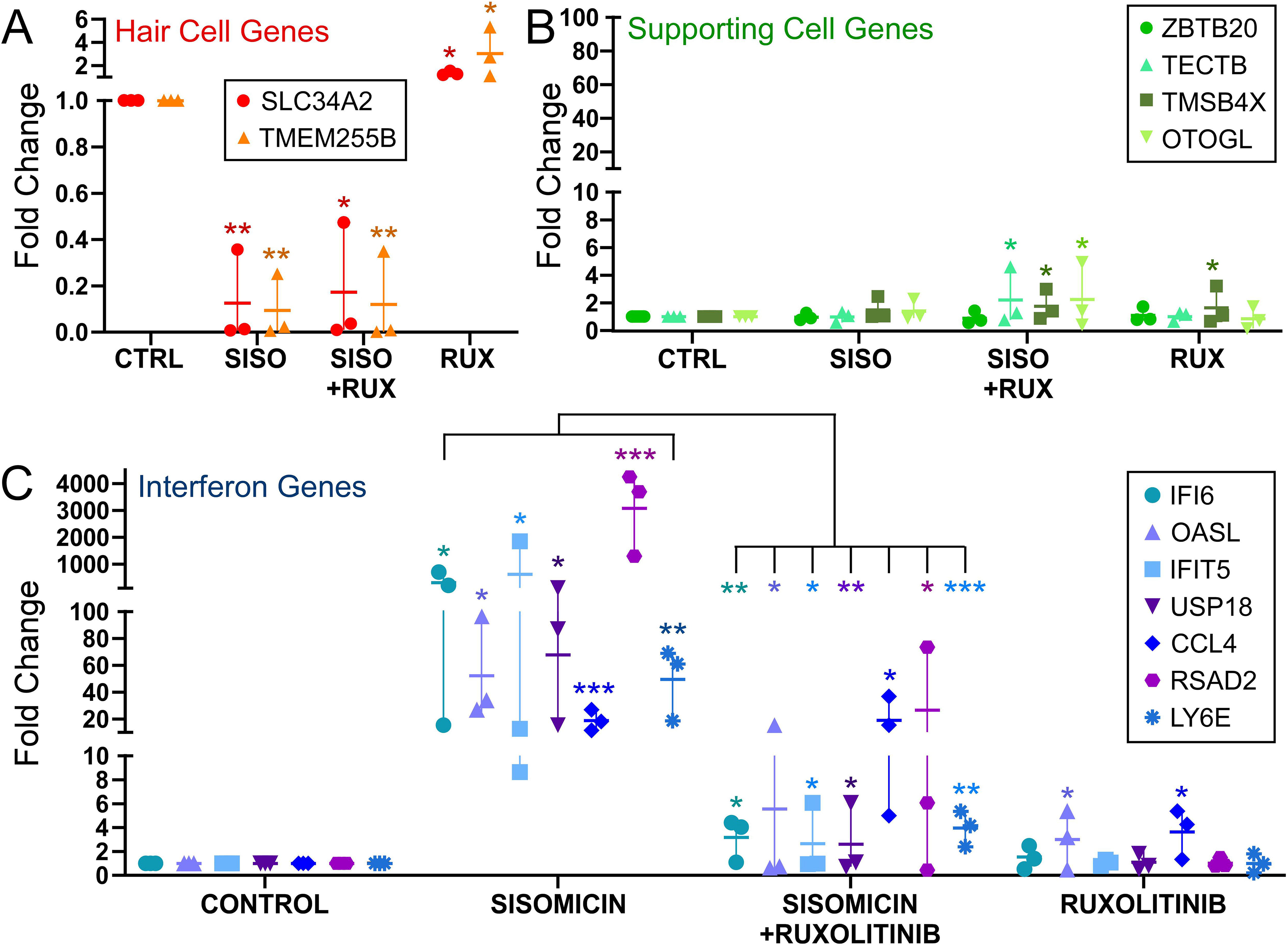
JAK/STAT signaling is required for the interferon response in culture. Peeled sensory epithelia from P7 chickens were cultured according to Burns et al., 2008, with some modifications described in Methods. The epithelia were incubated overnight +/− sisomicin and +/− JAK/STAT inhibitor (RUX = ruxolitinib; NIF = nifuroxazide). The following day, sisomicin was removed and the epithelia were cultured another day +/− JAK/STAT inhibitors. Gene expression was determined by the 2^−ΔΔCT^ method using *HSPA8* as the reference gene. Data are reported as fold induction over controls, representing three biological replicates, and the average expression indicated by the horizonal line. Statistics were conducted using repeated-measures one-way ANOVA with Bonferroni post-hoc analysis. All post-hoc comparisons are relative to control. In addition, comparisons between Sisomicin and Sisomicin + Ruxolitinib are presented for interferon genes. * = p ≤ 0.05; ** = p ≤ 0.01; *** = p ≤ 0.001.

### Regenerating Hair Cells Serve as a Streamlined Model for Hair Cell Development

Functional regeneration in the chicken basilar papilla requires new hair cell generation as well as the re-establishment of hair bundles, proper epithelial cell polarity, tonotopic location-specific hair bundle shape, and physiological specializations related to mechanotransduction and frequency tuning (Fettiplace and Kim, 2014). Moreover, new hair cells need to be rewired with efferent neurites, re-establish afferent auditory nerve connections, and adhere to the tectorial membrane. The time course of these specific processes and whether they occur independently has not been deeply investigated. We observed that 96 hours after sisomicin damage, new hair cells expressed the hair cell marker MYO7A but were still immature. They lacked bundles, were connected to the basement membrane via cytocaud processes, and expressed both the supporting cell marker SOX2 and the young hair cell marker TUBB3 (Benkafadar et al., 2021; Janesick et al., 2021a). In contrast, three weeks after sisomicin infusion, we could not detect histological differences between controls and sisomicin-infused specimens (**Fig 4E, F, F’**), suggesting that the regenerative processes have largely concluded. This agrees with previous reports based on anatomical data, although full functional recovery likely requires 4-5 months (Duckert and Rubel, 1993; Girod et al., 1991; Tucci and Rubel, 1990).

A new population of 32 hair cells is unequivocally detectable at 96 hours post-sisomicin infusion in our single cell RNA-seq dataset (**Fig. 2A, E**; State S1 in **Fig. 2B**). These hair cells robustly express *ATOH1* (**Fig. 2E**), a critical transcription factor in hair cell development (Bermingham, 1999). Because the regenerating chicken basilar papilla is mainly concerned with building new hair cells, but overall morphogenesis is complete, we consider our model to be a novel paradigm in which hair cell generation is independent from the complexities of embryonic organ development. We conducted differential gene expression analysis on the new hair cell cluster (State S1 in **Fig. 2B**), comparing its transcriptome to mature, non-compromised control hair cells (States S4∪S5 in **Fig. 2B**). This comparison revealed 851 higher expressed and 404 lower expressed genes (**Fig. 6A** and **Table S6**) using an FDR cut-off of 0.01 and | log_2_FC | > 2 in nascent hair cells. We concluded that new hair cells are transcriptionally distinct from mature hair cells.

**Figure 6.**
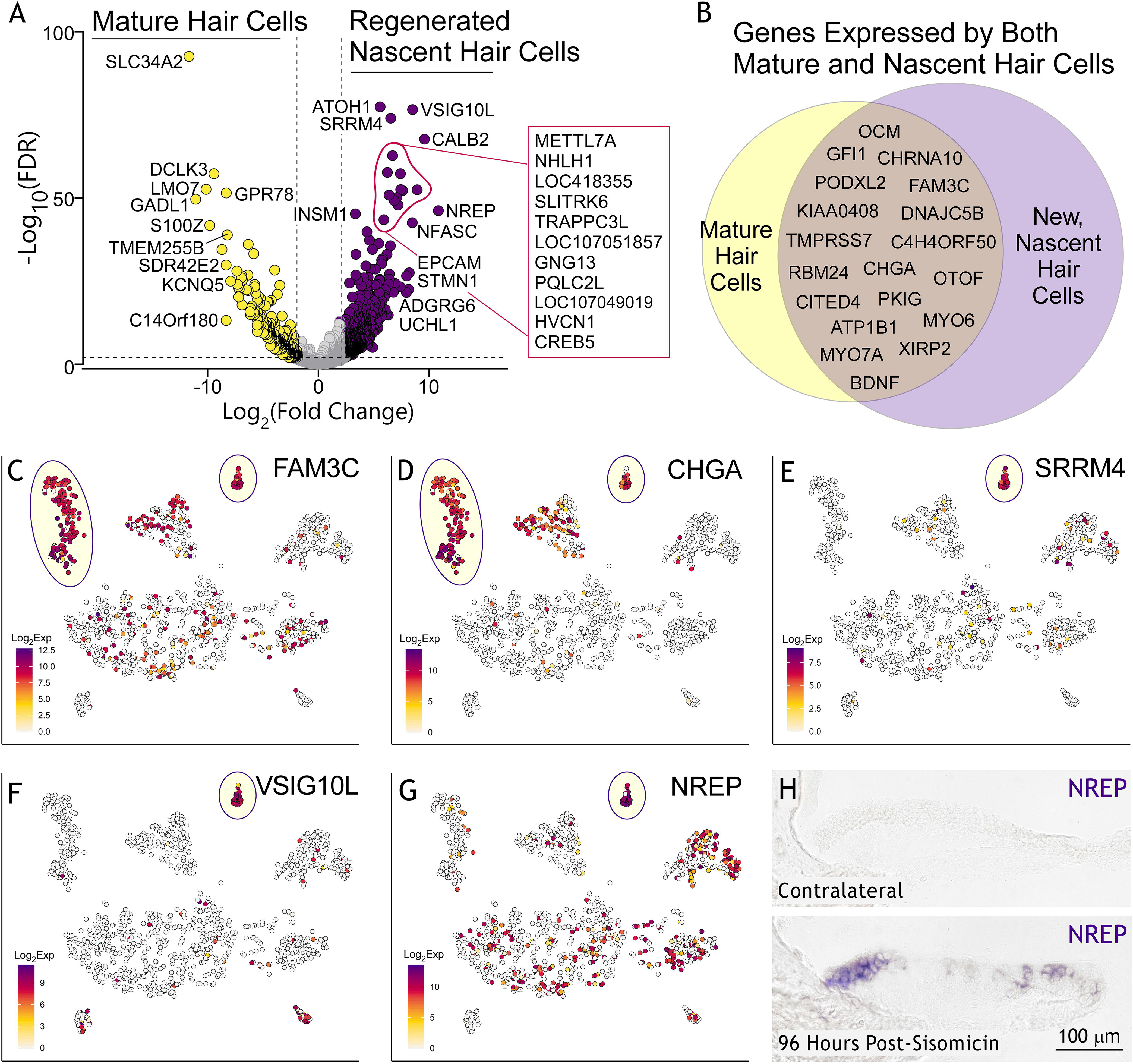
Novel markers of transcriptomally distinct new, regenerated hair cells. **(A)** Volcano plot illustrating genes expressed in new, regenerating hair cells (purple dots) at least 4-fold higher and significantly different (FDR < 0.01) compared with genes expressed in mature hair cells (yellow dots). Circled cells have gene labels listed in a box to the side of the volcano to avoid crowding. **(B)** Venn diagram comparing statistically significant new hair cell markers and mature hair cell markers. **(C-D)** tSNE plots project log_2_ transformed, normalized expression counts for *FAM3C* and *CHGA*. These are two novel genes that are representative of the Venn diagram overlapping region in (B). **(E-G)** tSNE plots for *SRRM4*, *VSIG10L*, and *NREP* which are representative genes of the new, regenerating hair cell cluster. Scalebar = log_2_ transformed, normalized expression counts. **(H)***In situ* hybridization validates *NREP* mRNA expression in new hair cells in P11 transverse sections, 96 hours post-sisomicin damage.

The present dataset provides a resource for identifying novel genes involved in hair cell differentiation that immediately precede or follow *ATOH1* expression. **Fig. 6A** and **Table S6** present genes that are exclusively expressed by new hair cells. A smaller collection of genes listed in **Fig. 6B** is expressed in both new and mature hair cells. This group includes well-characterized hair cell genes such as *MYO6*, *GFI1*, and *OTOF*, and novel genes such as *FAM3C* and *CHGA* (**Fig. 6C, D**). We highlight regenerating hair cell-specific genes *VSIG10L*, *SRRM4*, and *NREP* in **Fig. 6E-G**. We validated *NREP* mRNA in new hair cells at 96 hours post-sisomicin damage (**Fig. 6H**).

The chicken basilar papilla comprises three types of hair cells: superior tall, tall, and short (Janesick et al., 2021b). Tall and short hair cells may be analogous to mammalian inner and outer hair cells, respectively. Differential expression analysis revealed that most distinguishing tall and short hair cell markers are absent from new hair cells (**Fig. 6A, Table S6**). Two exceptions are *CXCL14*, a tall hair cell marker (Janesick et al., 2021b), and *INSM1*, an early mammalian outer hair cell marker (Lorenzen et al., 2015). *CXCL14* was present in embryonic day 11 chicken basilar papilla (Gordon et al., 2011), specifically in hair cells (**Fig. S5A**); we conclude that *CXCL14* is expressed in all hair cells during development and later becomes restricted to tall hair cells. We compared our top-scoring new hair cell markers using the gene Expression Analysis Resource (gEAR; Orvis et al., 2021; https://umgear.org/p?l=f7baf4ea) to genes expressed in *ATOH1*-positive neonatal mouse cochlear hair cells (Kolla et al., 2020; Orvis et al., 2021) and identified 14 genes conserved between chicken and mouse (**Fig. S6**).

The 96h post-sisomicin new hair cell cluster (State S1 in **Fig. 2B**) is the only group expressing *ATOH1* (**Fig. 2E**), which is expected because the P7 chicken basilar papilla is quiescent and does not yield *ATOH1* expression or EdU positive cells without damage. In contrast, the P7 utricle exhibits a low level of proliferation and hair cell turnover and transitory cells can be detected by their expression of *ATOH1* (Cafaro et al., 2007; Scheibinger et al., 2018). We compared the 96-hour dataset with chicken utricle sensory epithelial cells (Scheibinger et al., 2021a; https://umgear.org/p?l=bca63f4b; see Supplemental Files for preprint) using our analysis pipeline for filtering and normalization as described above. After quality control, 845 cells were examined with CellTrails (Ellwanger et al., 2018) representing three experiments: P11 control basilar papilla, P11 (96h post-sisomicin) sisomicin-infused basilar papilla, and the P7 utricle (**Fig. 7A**). We leveraged the information from Janesick et al., 2021b and Scheibinger et al., 2021 to identify the major cell clusters that appeared after dimension reduction (**Fig. 7B**).

**Figure 7.**
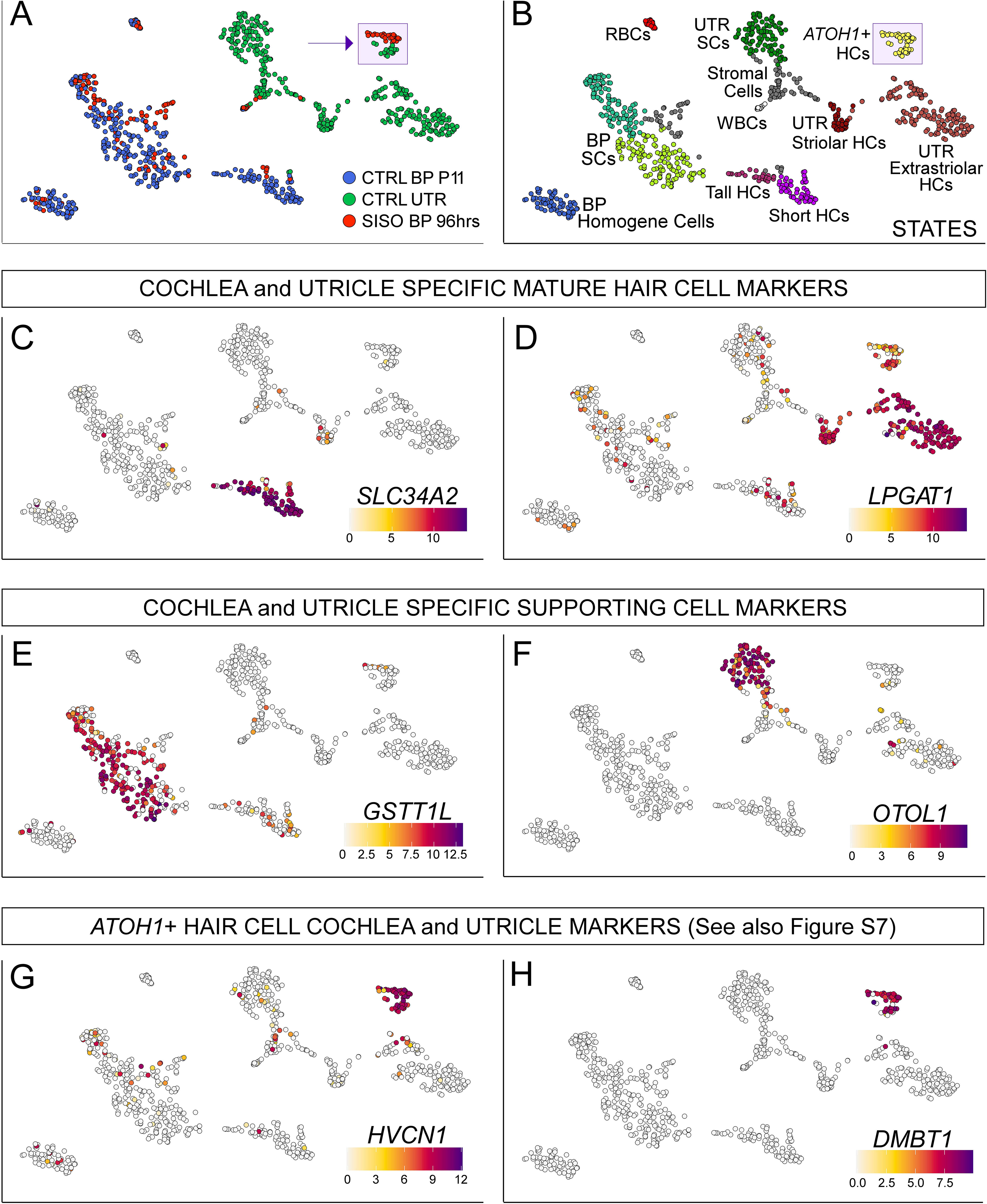
New basilar papilla hair cells and transitional utricular hair cells are highly resemblant and express analogous markers. **(A)** T-distributed stochastic neighbor embedding (tSNE) plots and CellTrails clustering of basilar papilla P11 and 96-hour post-sisomicin (SISO) cells in comparison to baseline P7 utricle cells. **(B)** All cell groups were identified using markers from Janesick et al., 2021 (baseline basilar papilla) and Scheibinger et al., 2021 (baseline utricle). HCs = hair cells; SCs = supporting cells; RBCs = red blood cells; WBCs = white blood cells; UTR = utricle; BP = basilar papilla. **(B, C)** New hair cells were identified by the expression of *ATOH1*. Basilar papilla and utricle *ATOH1*-positive cells assemble into a single cluster of the same CellTrails state. **(D-J)** Marker genes expressed by both new, regenerating basilar papilla hair cells and transitional utricular hair cells are projected onto the tSNE plots. Scalebar = log_2_ transformed, normalized expression counts. Note that *DMBT1* is known by two different names in our dataset: LOC112532717 and LOC107049019.

In mammals, utricle and cochlear hair cells are transcriptionally distinguishable as early as P1 (Burns et al., 2015). Our single-cell analysis confirmed that P7 avian utricle and basilar papilla supporting cells and hair cells also segregated from each other (**Fig. 7A, B**). The chicken basilar papilla epithelium expresses the specific marker *CALB1*, the hair cell marker *SLC34A2*, and supporting cell markers *GSTT1L* and *FGFR3*, exclusively (**Fig. 7C, E; Table S7, S8**). The chicken utricle expresses the hair cell-specific markers *CIB3*, *LPGAT1*, and *CRABP1*, and supporting cell-specific markers *HES5L* (*LOC419390*) and *OTOL1* (**Fig. 7D, F; Table S7, S8**). Regenerating new hair cells of the damaged basilar papilla and new transitory/turnover hair cells of the utricle cluster together and express *ATOH1* (**Fig. 7A, B; Fig. S7A-C**). We found that a large group of 2255 genes are shared between utricle and basilar papilla *ATOH1*-positive cells (**Fig 7G, H, Fig. S7C-H, Table S9**). We conclude that there is a universal early transcriptomal program for both regenerating cells of the basilar papilla and transitory cells of the utricle. Subsequently, these immature cells differentiate into more specialized vestibular and auditory mature hair cell fates. This concept was previously reported for *in vitro*-generated embryonic stem cell-derived nascent hair cells, based on functional assessments (Oshima et al., 2010).

### CALB2, USP18, and TRIM25 link responding supporting cells to new hair cells

Our next goal was to link the responding supporting cell population to new hair cells. The activation of the nascent hair cell transcriptomal program occurs between 38 and 96 hours post-sisomicin damage. The responding supporting cells at 30 and 38 hours (S9 in **Fig. 2B**, **Fig. 3**) were distinct from the nascent regenerated hair cells at 96 hours (S1 in **Fig. 2B**, **Fig. 6**). This gap exposed a limitation of our study where we would need to acquire additional cells from time points between 38 and 96 hours to enable the building of a proper trajectory between responding supporting cells and new hair cells. In lieu of a trajectory analysis, we compared the top-ranking genes of the new hair cell and the responding supporting cell group and found three genes present in both: *CALB2*, *USP18*, and *TRIM25* (**Fig. 8A-D**). These genes link the responding supporting cell population with the newly regenerated nascent hair cells and represent “cornerstones” of the trajectory of presumed gene expression changes that define the conversion of responding supporting cells towards new hair cells. We confirmed *USP18* and *CALB2* expression by *in situ* hybridization at 48 and 96 hours post-sisomicin, respectively (**Fig. 8B’, D’**). In mammals, *USP18* competes with JAK, thus preventing phosphorylation of downstream substrates, including STATs (Malakhova et al., 2006; Wilmes et al., 2015). We hypothesized that *USP18* acts by suppressing the interferon response, allowing regenerated hair cell differentiation to commence. Furthermore, *USP18* is expressed scarcely in *ATOH1*+ hair cells in the undamaged utricle, labeling new hair cells generated during natural turnover. Therefore, because it is linked to interferon genes, *USP18* is likely specific to responding supporting cells and regenerating hair cells and is not expressed when hair cells are naturally produced in the utricle without induced damage.

**Figure 8.**
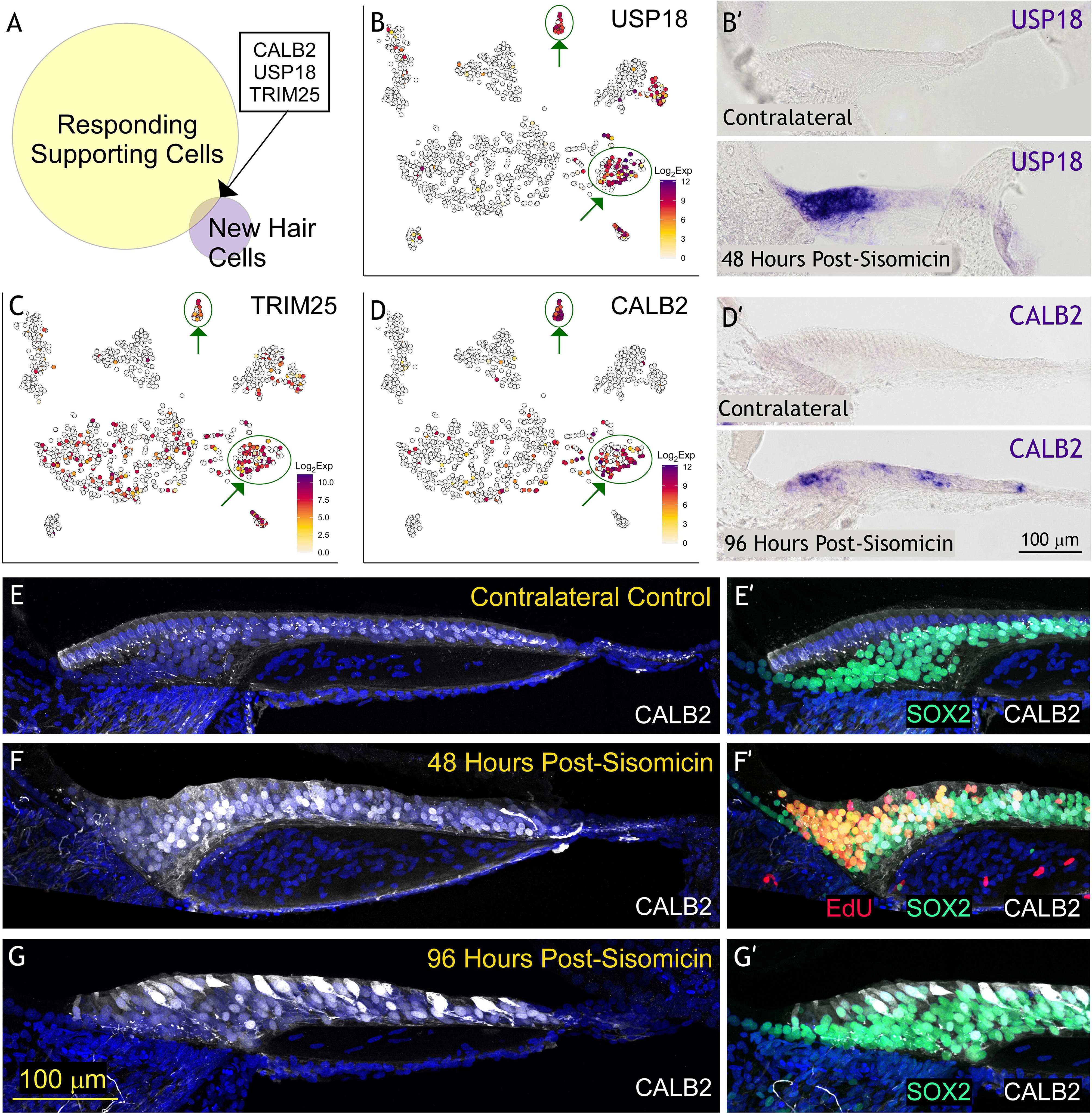
An exclusive set of genes associate with both responding supporting cells and new hair cells. **(A)** Venn diagram comparing statistically significant new hair cell markers and responding supporting cell markers. (B-D) tSNE plots project log_2_ transformed, normalized expression counts for the only three genes (*USP18*, *TRIM25*, and *CALB2*) that overlap between these two distinct clusters. **(B’)***In situ* hybridization validates *USP18* mRNA expression in responding supporting cells P9 transverse sections, 48 hours post-sisomicin damage. (**D’)***In situ* hybridization validates *CALB2* mRNA expression in new hair cells P11 transverse sections, 96 hours post-sisomicin damage. **(E-G)** Immunohistochemistry for CALB2 protein (also known as Calretinin) in white and DAPI in blue at the timepoints listed. **(E’-G’)** Medial region of the same sections in E-G showing SOX2 (supporting cells) in green and Calretinin in white. **(F’)** EdU was subcutaneously injected 2x into chickens at approximately 42 and 48 hours post-sisomicin, and were sacrificed at 54 hours post-sisomicin. Proliferative cells are labeled in red. Note that EdU staining was not performed on Control **(E’)** or 96 hours **(H’)** specimens because we have never observed proliferation here (Janesick et al., 2021a).

We explored *CALB2* in greater depth because it is expressed in both responding supporting cells and new hair cells and also because an antibody against human recombinant CALB2 (a.k.a. Calretinin) detects CALB2 in the hair cells of frog saccule (Edmonds et al., 2000), mammalian inner ear (Dechesne et al., 1994), chicken utricle (Ellwanger et al., 2018), and the chicken basilar papilla (**Fig. 8E-G’**). We detected CALB2 protein in supporting cells and hair cells. CALB2 is expressed immediately after proliferation ends in the rat utricle (Zheng and Gao, 1997) in mature extrastriolar hair cells (Ellwanger et al., 2018). We detected CALB2 expression as early as embryonic day 9 in the chicken basilar papilla, (**Fig. S5B**), prior to the onset of OTOF expression in hair cells at E11 (**Fig. S5C-D**). Therefore, CALB2 functions as both a developmental and mature hair cell gene in different contexts, in addition to its well-known role in sensory ganglia neuronal subtypes specification (Shrestha et al., 2018; Sun et al., 2018).

We evaluated CALB2 protein expression in regenerating and control sensory epithelia (**Fig. 8E-G’**). In contralateral controls, CALB2 was present mainly in synaptic terminals appearing as punctate staining, and weakly in supporting cell nuclei (**Fig. 8E, E’**). At 48 hours post-sisomicin infusion, CALB2 was found in both medial and lateral responding supporting cells (CALB2/SOX2+), with a strong signal in the nuclei (**Fig. 8F, F’**). EdU staining in the same specimen confirmed that these supporting cells were in the proliferative phase of regeneration (**Fig. 8F’**). At 96 hours post-sisomicin damage, CALB2 was found in supporting cells largely in the nuclei and in new hair cells where the protein is cytoplasmic and nuclear (**Fig. 8G, G’**). It was previously known that CALB2 is an important marker of differentiated hair cells with functional physiological roles as a calcium sensor (reviewed in Camp and Wijesinghe, 2009). Here we showed that CALB2 can also be detected in responding and proliferating supporting cells in the chicken basilar papilla after damage and that the protein is abundant in regenerated nascent hair cells.

## Discussion

### Hair cell loss induced an interferon response in avian otic epithelial supporting cells

How non-mammalian species restore hearing after damage is among the most inspirational unanswered questions in the regeneration field. To address this question, we performed an RNA-seq analysis of the regenerating chicken basilar papilla. We collected cells at 48 hours post-damage for bulk RNA-seq and at 30-, 38- and 96-hours post-damage for single-cell RNA-seq. Our results revealed a striking induction of interferon response genes in supporting cells 30, 38, and 48 hours after sisomicin damage.

Inflammation and oxidative stress responses to ototoxic insult have been reported (reviewed in Dinh et al., 2015), and infiltrating leukocytes are often implicated in these responses (Bhave et al., 1998; Hirose et al., 2017; Kaur et al., 2015; Ladrech et al., 2007; O’Halloran and Oesterle, 2004; Tornabene et al., 2006; Warchol, 1997; Warchol et al., 2012). Other results demonstrated that supporting cells emulate immune cells and are “glial-like,” expressing GFAP and GLAST (Hayashi et al., 2020; Matsunaga et al., 2020; Monzack and Cunningham, 2013; Wan et al., 2013). Our baseline analysis of the chicken basilar papilla demonstrated strong expression of the astrocytic marker *ZBTB20* in supporting cells (Janesick et al., 2021b; Nagao et al., 2016). Therefore, supporting cells might already be primed to participate in neuroimmune responses.

Because previous reports investigating gene expression changes following hair cell damage in the chicken inner ear did not include single-cell transcriptome analysis, it was challenging to use these data to identify which cell types expressed immune-response genes (**Table S1**). Here we showed that damage induces strong upregulation of interferon response genes in supporting cells without significant contribution from blood cell infiltration. This observation is supported by a recent study using cultured streptomycin-treated chicken basilar papillae (Matsunaga et al., 2020). Our analysis focused solely on the peeled sensory epithelium so it is possible, albeit unknown whether interferon signaling might emanate from stromal or strial compartments of the basilar papilla. Interferon receptors (Goossens et al., 2013) are not expressed in the basilar or utricular sensory epithelium in our single-cell datasets (Janesick et al., 2021b; Scheibinger et al., 2021a). Therefore, the interferon response may be triggered by a different mechanism. A recent review article characterized all pattern-recognition receptors used by avian species to invoke an interferon response (Neerukonda and Katneni, 2020). We compared these immune sensors with our single-cell datasets but did not find any strong leads. The RNA-sensors, TLR3 and OASL were induced after damage but are expressed in supporting cells at homeostasis. Regardless of the initial trigger, the interferon response is typically communicated via JAK/STAT signaling, although other non-canonical pathways (MAPK, mTOR, pI3-kinase) can contribute to the response. We found that JAK/STAT signaling was required because Ruxolitinib could effectively block expression of the damage-induced interferon genes.

Like all epithelial cells, supporting cells function as a barrier to the external environment. After hair cell death, supporting cells expand their apical surfaces to compensate for lost hair cells (reviewed in Francis and Cunningham, 2017). This controlled process preserves epithelial integrity and endocochlear potential. In other epithelial systems, there is ample evidence that cells *not* from the hematopoietic lineage can provide a barrier to the external environment and mimic some of the actions of immune cells. For example, epidermal keratinocytes and dermal fibroblasts can secrete interleukins, apolipoproteins, and antimicrobial peptides (reviewed in Nguyen and Soulika, 2019). Zebrafish ectoderm-derived “metaphocytes” share transcriptomic and morphological characteristics with macrophages (Lin et al., 2019). These metaphocytes sample soluble antigens directly from the external environment and transfer them to macrophages, but their developmental lineage is different from the macrophages they service (epidermis *versus* mesodermal germ layer).

Whether the upregulation of interferon response genes that we observe is a trigger for regeneration and repair or rather a mechanism to mitigate the effects of damaged cells is currently unknown and the subject of future study. In other systems, interferon signaling is linked to epithelial repair and also to cell turnover and proliferation. After a colonic injury, the intestinal epithelium mounts an interferon response that is required for regeneration and depends on EGFR and the ligand AREG (McElrath et al., 2021). Mice lacking the interferon receptors show impaired epithelial proliferation, compromising the mucosal barrier (McElrath et al., 2021). Chronic viral infection also promotes epithelial proliferation in the kidney, liver, salivary glands via type I interferon signaling (Sun et al., 2015). In the *Drosophila* midgut, an injury-induced cytokine response promotes stem cell and enterocyte division and differentiation, respectively (Jiang et al., 2009). Therefore, there is support in multiple models for interferon responses leading to S-phase re-entry and regeneration.

### Controlling the Interferon Gene Response in the Chicken Basilar Papilla

As H.W. Longfellow wrote, “Great is the art of beginning, but greater is the art of ending.” Signal inactivation mechanisms are built into developmental and regenerative systems to prevent runaway feed-forward signaling. Unregulated interferon signaling caused neuronal damage by impairing neurotrophic activity (Dedoni et al., 2012). The chicken basilar papilla appears to have excellent control over the interferon response. Despite how massively as *IFI6*, *IFIT5*, and *OASL* (among other genes) are induced at 48 hours post sisomicin, their expression is almost non-existent in regenerated nascent hair cells. We hypothesized that *USP18* could play a key role in shutting down interferon response genes, considering its known role in terminating the interferon signaling cascade (Basters et al., 2018; Hou et al., 2021). *USP18* is one of few genes that is expressed in both supporting cells at 30 and 38 hours (FC = 15, FDR = 3.1 × 10^−51^), as well as nascent hair cells at 96-hours post-sisomicin (FC = 60, FDR = 9.0 × 10^−21^). *USP18* also appears to be specific to injury since we did not find it expressed in transitional utricle cells that turn over naturally rather than in response to injury. Another negative regulator of JAK/STAT signaling, *SOCS3*, is sporadically expressed in new hair cells (FC = 6.3, FDR = 1.7 × 10^−11^) and is also important for hair cell regeneration in zebrafish after noise damage (Liang et al., 2012).

*TRIM25* is another gene that straddles two groups in our dataset: responding supporting cells (FC = 5.1, FDR = 3.8 × 10^−10^) and new hair cells (FC = 12.6, FDR = 2.4 × 10^−14^). *TRIM25* is an interferon-inducible E3 ubiquitin ligase that participates in the innate immune response, regulation of cell proliferation, as well as cancer cell invasion (reviewed in Martín-Vicente et al., 2017). TRIM25 is an important cofactor in LIN28-mediated uridylation of pre-let-7 (Choudhury et al., 2014). Intriguingly, LIN28B is expressed in the mammalian cochlea and controls the ability of neonatal murine auditory supporting cells to generate hair cells through mTOR signaling (Li and Doetzlhofer, 2020). Neither LIN28A nor LIN28B were expressed in the chicken basilar papilla by single-cell RNA-seq. Hence, this mechanism in mammals to generate hair cells is either not conserved or utilizes different gene sets in avian species. TRIM25 positively regulates interferon signaling through K63-linked ubiquitination of RIG-I, but also negatively regulates interferons via stabilization of FAT10, and the ubiquitination of MAVS (reviewed in Martín-Vicente et al., 2017). The RIG-I homolog is absent in chickens (Santhakumar et al., 2017), thus likely altering TRIM25 functionality in comparison to mammals, in the context of positively or negatively regulating interferons.

In various non-regenerating adult mammalian tissues, an inflammatory response contributes to fibrosis and the excess production of collagen and other extracellular matrix proteins. When the mammalian spinal cord is injured, a glial scar forms, impairing neuroplasticity, axon outgrowth, and regeneration (Alizadeh et al., 2019; Tran et al., 2018). Cardiac fibrosis after myocardial infarction impairs regenerative capacity in the adult mammalian heart (Talman and Ruskoaho, 2016). After damage, the mouse inner ear forms a phalangeal scar tissue marked by an epithelial-to-mesenchymal transition, cell migration/remodeling, and de-specialization of supporting cell subtypes (Lee et al., 2021). Compared to organ of Corti supporting cells, avian supporting cells are not particularly specialized to begin with. While they undergo extensive shape changes and a partial mesenchymal-to-epithelial transition (Hu and Corwin, 2007), scarring does not occur (reviewed in Stone and Rubel, 2000). *GFAP* is a significant contributor to glial scars, which inhibit neuronal regeneration following brain or spinal cord injury (Eddleston and Mucke, 1993; Hatten et al., 1991). The P15 mammalian cochlea expresses *GFAP* in inner border and inner phalangeal cells and Deiters’ cells (Rio et al., 2002). This expression persists in the aminoglycoside-damaged, flat epithelium (Ladrech et al., 2017). Intriguingly, our single-cell data revealed no expression of *GFAP* in supporting cells at baseline or after damage, thus exposing a potentially key difference between mammalian and avian damage responses.

### CALB2 is expressed in responding supporting cells after damage

*CALB2* also belongs to the small group of genes expressed in both responding supporting cells (FC = 12, FDR = 1.9 × 10^−29^) and new hair cells (FC = 745, FDR = 2.1 × 10^−68^) in our dataset. Calretinin is a calcium sensor expressed in distinct neuronal populations innervating sensory organs, including retinal ganglion cells, the granular layer of the cerebellar cortex, and brainstem auditory neurons (reviewed in Camp and Wijesinghe, 2009). This Ca^+2^-binding protein modulates neuronal excitability by functioning as a slow calcium chelator at resting intracellular Ca^+2^ levels and a fast-onset buffer at elevated intracellular Ca^+2^ levels (Gall et al., 2003; Schwaller, 2009). Calretinin is dynamically expressed during development with varied subcellular localization. For example, in the organ of Corti, calretinin is initially expressed in both inner and outer hair cells but disappears in outer hair cells upon maturity, coinciding with the loss of afferents (Dechesne et al., 1994). In the turtle auditory papilla, nuclear labeling of calretinin predominates over cytoplasmic localization (Hackney et al., 2003). In the adult chicken basilar papilla, we found calretinin expression in the puncta of nerve terminals associated with tall hair cells and observed crescent-shaped staining around short hair cells, resembling a calyx. We also detected calretinin in the body of superior tall hair cells and faint nuclear staining in supporting cells.

Forty-eight hours after damage, nuclear expression of CALB2 in supporting cells intensified and persisted as newly regenerated CALB2+ hair cells emerged. To our knowledge, this is the first time CALB2 was observed strongly in proliferating supporting cells during development or regeneration. In the rat utricle, CALB2 was observed only in post-mitotic, differentiating hair cells (Zheng and Gao, 1997). In the chicken utricle, CALB2 was detected in new hair cells of an asymmetric pair but not in the supporting cell layer during both natural turnover and regeneration (Stone and Rubel, 1999). Therefore, the supporting cell expression of CALB2 is unique to the regenerating chicken basilar papilla. The nuclear staining we observed for CALB2 is also unusual, although there is precedent for nuclear labeling in turtle hair cells (Hackney et al., 2003), as well as nuclear translocation of calretinin in human colon carcinoma cells in response to vitamin D (Schwaller and Herrmann, 1997). The Vitamin D receptor is not expressed in the chicken basilar papilla sensory epithelium; therefore, it is unlikely that calretinin subcellular localization is regulated via the vitamin D receptor. Whether hair cell differentiation requires early, nuclear CALB2 localization is unknown. In the mammalian inner ear, CALB2 expression in new hair cells requires ATOH1 (Bermingham et al., 1999). It is unlikely that ATOH1 would require CALB2 expression in hair cell development, despite that we observe an abundance of CALB2 mRNA and protein before new hair cells emergence and before *ATOH1* expression. Rather, we infer that CALB2 expression could reflect unique calcium needs of damaged supporting cells. Indeed, interferon signaling has been known to increase calcium influx in microglial cells (Franciosi et al., 2002).

### Regenerative Strategies Inferred at Three Weeks Post-Sisomicin Damage

There are two modes of hair cell regeneration in the chicken sensory epithelium: proliferation and phenotypic conversion. Hair cells on the medial/neural side of the epithelium incorporate thymidine analogs after damage (reviewed in Janesick and Heller, 2019). In contrast, hair cells on the lateral/abneural side transdifferentiate from a supporting cell to a hair cell directly, rather than undergoing a mitotic event (reviewed in Janesick and Heller, 2019). Previous studies focused on the chicken basilar papilla six days post-gentamicin and found that 34% of newly generated short hair cells incorporated bromodeoxyuridine, compared to 81% of new tall hair cells (Cafaro et al., 2007). Because the time course of regeneration is on the order of months (Duckert and Rubel, 1993; Girod et al., 1991; Tucci and Rubel, 1990), these studies represent only an early snapshot before the full complement of hair cells are restored. We extended our analysis to three weeks post-sisomicin and found that hair cells on the medial/neural side of the epithelium were regenerated predominantly by mitotic events (87%). Only 21% of new lateral hair cells incorporated EdU.

In the vestibular system, supporting cell replenishment was postulated as a mechanism to replace supporting cells that had directly converted to hair cells without dividing (Scheibinger et al., 2018). However, we found that only 4% of lateral supporting cells proliferated. For example, across all sections, we quantified on average 28 EdU-negative short hair cells but only 13 EdU-positive supporting cells beneath them. We envision two alternative conclusions from this result. The first is that replenishment of supporting cells was simply incomplete or not required. The ratio of lateral supporting cells to short hair cells is over twice the ratio of medial supporting cells to tall hair cells (Goodyear and Richardson, 1997; Janesick and Heller, 2019), suggesting that an abundant reservoir of supporting cells is available. It would be informative to investigate whether the lateral supporting cells could eventually be depleted after repeated rounds of damage and regeneration. The second conclusion is that our window of EdU administration was too early/short and might not have detected a second/later wave of supporting cell division. A third conclusion is that medial supporting cells could migrate to the lateral side over time, but the time course is too gradual/slow to observe. Indeed, we found that 3.4x more medial supporting cells divided than did tall hair cells, suggesting that supporting cells are not simply replacing themselves with asymmetric divisions, but might undergo symmetric divisions as well, and then potentially migrate to the lateral side.

The possibility that regeneration of the sensory epithelium is incomplete (both at the hair cell and supporting cell level) has merit because previous reports have observed that functional hearing is not entirely restored. In pigeons and chickens, aminoglycoside damage triggers a regenerative response that eventually yields substantial hearing recovery. However, high-frequency regions (>1.5kH) still exhibit a 20-30 decibel permanent threshold shift at 4-5 months post-damage (reviewed in Dooling et al., 2008; Saunders and Salvi, 2008). We observed that the timescale of short hair cell replenishment is slower than for tall hair cells. The density of tall hair cells at three weeks approximates control levels, whereas the short hair cell regions are more sparsely populated. If tall hair cells fulfill a similar role to mammalian inner hair cells, then it would make sense to prioritize tall hair cell restoration (hearing function) over short hair cells (amplification).

### Nascent Hair Cells are Incipient Modified Neurons

The generation of sensory hair cells within the inner ear during regeneration and development relies on the expression and regulation of early transcription factors such as ATOH1, PROX1, and GFI1. Manipulation of these genes can produce hair cell-like cells that often fail to reach full maturity (Gubbels et al., 2008; Menendez et al., 2020; Oshima et al., 2010). This could be an artefact of culture conditions or could reflect the absence of other key players in hair cell development. Misexpression of *ATOH1* mRNA in the developing mouse cochlea produces functional hair cells (Costa and Henrique, 2015; Menendez et al., 2020; Sayyid et al., 2019), suggesting that competence for reprogramming exists in embryonic cochlear progenitor cells. Our 96-hour timepoint revealed a distinct population of young hair cells that express novel markers likely to precede or follow ATOH1 expression. We anticipate that these markers will provide a valuable resource for placing ATOH1 expression into a cascade of transiently expressed accessory genes required for the generation of *bona fide* hair cells.

The most striking similarity among genes expressed in nascent hair cells is that many belong to the biological process of neurite outgrowth and projection. Hair cells can be categorized as compact neurons (Heller et al., 2002) and use chemoattractive signals such as BDNF to establish and maintain connections with afferent nerve cells. After hair cell loss, the peripheral neural processes retract, but remain near the basement membrane. It appears that new hair cells then “cue” the spiral ganglia for immediate rewiring. For example, *SRRM4* is a deafness gene that is important for neurogenesis, neurite outgrowth, and axon guidance (Calarco et al., 2009; Quesnel-Vallières et al., 2015). We found that *SRRM4* is 90-fold higher expressed (FDR = 1.1 × 10^−74^) in the 96-hour, new hair cell group compared to mature hair cells. *NFASC* (FC = 347, FDR = 3.2 × 10^−43^) is a cell adhesion molecule expressed in immature olfactory sensory neurons (McIntyre et al., 2010) and was recently linked to auditory neuropathy spectrum disorder (Harper et al., 2020). *SLITRK6* (FC = 169, FDR = 3.5 × 10^−53^) has structural similarity to NTRK neurotrophin receptors and was expressed in specific regions of the embryonic mouse thalamus and organ of Corti (Aruga and Mikoshiba, 2003; Brose and Tessier-Lavigne, 2000; Katayama et al., 2009). SLITRK6-deficient mice showed profound loss of cochlear innervation (Katayama et al., 2009), and mutations in humans cause neuropathy and deafness (Ordonez and Tekin, 2017). *NREP* (FC = 1786, FDR = 7.8 × 10^−47^) was found in the regenerating chicken basilar papilla (Hawkins et al., 2006) and in transitional cells of the utricle (Scheibinger et al., 2021a). NREP was up-regulated after axotomy and enhanced neurite genesis and elongation in facial nerve regeneration (Fujitani et al., 2004).

In further support of our model that new hair cells are burgeoning neurons, we also identified *NHLH1* (FC = 73, FDR = 2.4 × 10^−58^), a putative transcription factor that was expressed in early post-mitotic neurons of the fetal brain (Lipkowitz et al., 1992; Murdoch et al., 1999), and in sensory epithelia and ganglia of the retina, tongue, and skin (Krüger et al., 2006). NHLH1 was expressed in nascent mouse hair cells and developing vestibulocochlear ganglia, but not in mature hair cells (Jahan et al., 2010; Krüger et al., 2006). *DMBT1* (LOC107049019 in our dataset) is another neurogenesis gene expressed exclusively in new hair cells (FC = 20; FDR = 1.6 × 10^−40^). Mutations in DNMT1 caused hereditary sensory neuropathy and hearing loss (Ding et al., 2018; Klein et al., 2011).

There are ~20 genes shared between new, nascent hair cells and mature hair cells, including many known genes such as *OCM*, *GFI1*, *CHRNA10*, *OTOF*, *MYO6/7A*, *XIRP2*, and *BDNF*. We also identified several other genes that have not been well-characterized in the inner ear. FAM3C is a candidate gene for autosomal recessive nonsyndromic hearing loss locus 17 of unknown function (Pilipenko et al., 2004). CHRNA1 is predicted to be an early and direct target of ATOH1 (Scheffer et al., 2007). DLK2 (Delta Like Non-Canonical Notch Ligand 2) is an EGF-like protein that functions as an inhibitory non-canonical ligand of NOTCH1 receptors (Sánchez-Solana et al., 2011). CHGA is a pan-neuronal marker downstream of Notch signaling (Ratié et al., 2013; Ratié et al., 2014). Inventorying these novel markers contributes to the toolset for future experiments aimed at building mature and integrated hair cells in culture, from stem cells, or via *in vivo* reprogramming.

### Biophysical Properties of Mature Hair Cells Can be Inferred by Comparison to Young Hair Cells

Our dataset lends itself to distinguishing mature from newly regenerated hair cells. Identification of genes that are only present in mature hair cells provides information about requirements hair cells have after they mature. There are numerous hair cell genes not expressed by nascent hair cells, and we predict that these genes are involved in later processes related to hair bundle development, mechanotransduction, and hair cell subtype-specific specializations. We and others have observed small hair bundles by seven days after aminoglycoside damage (Duckert and Rubel, 1990; Janesick et al., 2021a). At 96-hours, the timepoint collected for this dataset, the hair cell is likely still in a primary differentiation phase and has not commenced bundle development and mechanotransduction. For example, *LMO7* is 1160-fold higher expressed (FDR = 2.7 × 10^−53^) in mature hair cells compared to the 96-hour, new hair cell group. *LMO7* is localized to the cuticular plate in mice and is responsible for F-actin and stereocilia organization (Du et al., 2019). Since cuticular plate development parallels stereocilia development (Duckert and Rubel, 1990), we argue that *LMO7* is a late-onset gene required only after hair cell commitment and differentiation has occurred around 72-96 hours post-sisomicin. Similarly, *DCLK3* (FC = 710, FDR = 5.8 × 10^−58^), which interacts with both tubulin and actin (Dijkmans et al., 2010), is likely only to be required when stereocilia development commences.

Nascent hair cells lack hair cell genes such as *KCNQ5* (a voltage-gated potassium channel), *S100Z* (a calcium-binding protein), and *SLC34A2* (a sodium-dependent phosphate transporter) (Gillespie and Müller, 2009). We previously identified *GADL1* as novel and abundantly expressed hair cell marker, and in this present study, we found it only in mature hair cells (FC = 2224, FDR = 2.5 × 10^−50^). *GADL1* decarboxylates amino acids and is important for the biosynthesis of taurine (Liu et al., 2012; Mahootchi et al., 2020), which modulates intracellular calcium homeostasis in the cochlea (Liu et al., 2006).

Nascent basilar papilla hair cells appear to express a common hair cell program and cannot be distinguished readily from transitional *ATOH1*-positive utricle cells. Likewise, such specializations in hair cells as “tall,” “short,” or “superior tall” (Janesick et al., 2021b) are probably not realized until later time points. In support of this, we observed two genes which we definitively assigned to the short hair cell identity, *C14orf180* (FC = 329, FDR = 8.0 × 10^−14^) and *SDR42E2* (FC = 330, FDR = 1.7 × 10^−30^), that were found exclusively in mature hair cells. However, it should be noted that *CXCL14*, a mature, “tall” hair cell marker (Janesick et al., 2021b), was expressed in the developing avian basilar papilla (Gordon et al., 2011), utricle hair cells (transitional and mature), and in new, regenerated hair cells (FC = 29, FDR = 9.8 × 10^−6^). It was previously observed that all embryonic basilar papilla hair cells have a slender morphology before the lateral hair cells take on a squattier, “short” appearance (Fischer, 1992; Munnamalai et al., 2017). We infer from these data that the default hair cell subtype is “tall.” On the other hand, we found that *INSM1*, an outer hair cell determinant in the mammalian cochlea (Lorenzen et al., 2015; Wiwatpanit et al., 2018), is modestly expressed in new hair cells (FC = 9.9, FDR = 6.8 × 10^−46^), and completely absent from mature hair cells. Since evidence for homology between short and outer and tall and inner markers is scarce (Janesick et al., 2021b), we can only conclude that *INSM1* is a hair cell development gene rather than a specific determinant of tall versus short fate.

## Acknowledgements

We thank Drs. T.A. Jan and D. Ellwanger for help with data analysis and visualization, Drs. Y. Song and R. Hertzano for expert assistance with gEAR, and members of the Heller laboratory for helpful discussions. Cell sorting/flow cytometry analysis for this project was conducted in the Stanford Shared FACS Facility. The Stanford Functional Genomics Facility (M.C. Yee, S. Sim, and V. Shokoohi) was utilized for processing of cells and sequencing. Imaging was done in the OHNS microscopy core, and we thank Drs. L. Becker, P. Atkinson, and M. Bartho for their excellent training and help with equipment. Funding was provided by the A.P. Giannini Foundation (AJ), School of Medicine Dean’s Postdoctoral Fellowship (AJ, NB), the Stanford Initiative to Cure Hearing Loss, and by the Hearing Restoration Project consortium of the Hearing Health Foundation.

## Materials and Methods

### Bulk (Population-Based) RNA-sequencing and analysis

All surgical procedures and dissections were performed according to Janesick et al., 2021a,b. Each epithelium was placed into 100 μL RNAqueous™ Lysis Solution in a nuclease-free tube, quickly triturated and vortexed, then frozen at −80°C. RNA was extracted using the RNAqueous™-Micro Total RNA Isolation Kit, following the manufacturer’s instructions. Total RNA was assessed with the Agilent 2100 Bioanalyzer at the Stanford Functional Genomics Facility (SFGF). The RNA yield from one peeled, sensory epithelium is approximately 20-30 ng with RIN values ranging from 8.2 to 9.5. Library preparation was performed by SFGF using the SMART-Seq® v4 Ultra® Low Input RNA Kit (Takara Bio Inc). The libraries were sequenced on the Illumina HiSeq 4000 platform (2 × 75 bp paired-end reads). N = 3 (for control) and N = 3 (for sisomicin), when 1N = 1 cold peeled epithelium. No pooling of samples was done, thus minimizing the averaging of biological variability.

Data from SFGF were provided as raw Fastq files which we submitted to BasePair (www.basepairtech.com). Reads were pre-processed with fastp v0.19.4 (Chen et al., 2018) and aligned with STAR v2.6 (Dobin et al., 2013) to Gallus gallus genome GRCg6a. BasePair provided gene feature counts (see **Table S10**) which we processed by following and combining the methods outlined in two software tool articles (Chen et al., 2016; Moisan et al., 2014). We filtered out 10806 of 22758 genes by employing the HTSFilter v3.8 (Rau et al., 2013) on trimmed-mean-of-M-values (TMM) normalized count data to yield a statistically robust, non-arbitrary threshold for low abundance and constant genes. We imported filtered genes into edgeR v2.14 for differential gene expression analysis (Robinson et al., 2010). Mean-difference volcano and per-sample expression plots were generated using Glimma (Su et al., 2017). For interactive Glimma plots, see Supplemental Files. The volcano plot in **Fig. 1** was manually colored using Canvas × Draw v20. Heatmap of log_2_(counts-per-million) values was created using Pheatmap v1.0.12. Gene ontology analysis was conducted using PathfindR v1.6.1 with the Reactome database (Jassal et al., 2019; Ulgen et al., 2019).

### Single Cell RNA-sequencing and analysis

All methods for single cell RNA-seq and data processing were conducted as described in Janesick et al., 2021b. The experimental design is outlined in **Fig. S2**. Cold-peeled epithelia from 4-6 animals belonging to one clutch of chickens were dissociated, washed, and submitted for FACS. Cells were index sorted into two 96-well plates (Batch 1&2) after excluding for debris, doublets, and low viability. This process was repeated twice more (per timepoint) with different clutches of chickens (Batch 3&4 and Batch 5&6). Cells were submitted to the Stanford Functional Genomics Facility where they were processed using the Smart-Seq2 protocol (Picelli et al., 2014). cDNA size distribution was assessed for each individual cell using an Agilent 2100 Bioanalyzer and 364 cells were selected for library construction. All cell libraries were sequenced together with an Illumina NextSeq 500 sequencer aiming for 400 million 150 bp, paired end reads resulting in about 1 million reads per cell.

Barcode demultiplexing and read mapping against the Gallus gallus genome (release GRCg6a; https://www.ncbi.nlm.nih.gov/assembly/GCF_000002315.6) was conducted as described in Janesick et al., 2021b. Quality control was conducted in R using scater v1.20.1 (McCarthy et al., 2017). Information-poor cells were removed based on the following thresholds: number of raw counts > 10,000; number of expressed genes > 100. Lowly-expressed genes were also removed (average counts > 0.1). Normalization was performed with the SCnorm package (Bacher et al., 2017). Non-linear dimensionality reduction and cell clustering were conducted using the CellTrails package (Ellwanger et al., 2018). Differential expression analysis between clusters was performed with EdgeR (Robinson et al., 2010) on log_2_ normalized expression counts. False discovery rates are reported as Benjamini and Hochberg corrected p-values. tSNE and volcano plots were generated as previously described (Janesick et al., 2021b). Violin plots were generated in ggplot2 v3.3.4.

### Data Availability and Preprints

The RNA sequencing data generated in this paper is available from GEO with accession number GEO: “TBD in progress”. The data is also available at gEAR, a gene Expression Analysis Resource (Orvis et al., 2021), via https://umgear.org/xxxxxxx. The normalized log count matrices and Single Cell Experiment containers are available on Zenodo (https://doi.org/xx.xxxx/zenodo.xxxxx). The preprints for Janesick & Scheibinger, 2021a and Scheibinger et al., 2021 have been uploaded as supplemental files. Interactive Glimma plots for visualizing and searching the bulk RNA-seq data are contained in a folder in Supplemental Files. Download the <u>entire</u> folder, unzip, and view “MD-Plot.html”.

### Vibratome sectioning, in situ hybridization, and immunohistochemistry

Basilar papillae ducts in bone were processed for vibratome sectioning and *in situ* hybridization or immunohistochemistry as previously described (Janesick et al., 2021b; Scheibinger et al., 2021b). T7 adapted RNA probe templates were prepared via PCR amplification with primers listed in **Table S11**. 5-ethynyl-2’-deoxyuridine (EdU) staining was conducted as previously described (Janesick et al., 2021a). Antibody sources for immunohistochemistry are provided in **Table S12**.

### Three Weeks Post-Sisomicin Experiment

Maintaining animals for longer than one week post-sisomicin requires extra attention to animal welfare due to vestibular defects from the sisomicin infusion (Janesick et al., 2021a). Furthermore, quantitation of EdU-positive cells must be conducted with a constant influx of EdU into the inner ear. Otherwise, an EdU-negative, MYO7A-positive new hair cell could be misconstrued for a phenotypically converted supporting cell, when instead, the cell simply did not incorporate EdU due to lack of EdU bioavailability. We determined that subcutaneously injected EdU will be available to the inner ear for approximately six hours (Janesick et al., 2021a). Some researchers choose a mini-osmotic for the constant delivery of thymidine analogs (Cafaro et al., 2007). We choose to re-administer 50 mg/kg EdU in 200 μL PBS/DMSO subcutaneously every 6 hours. We started dosing EdU at 30 hours post-sisomicin, and continued every 6 hours until the last dose at 80 hours (over 72 hours post-sisomicin). This dosing regimen ensured that we would cover the proliferative window (Janesick et al., 2021a).

We quantitated hair cells and supporting cells using a 3D virtual reality strategy outlined in Janesick et al., 2021a. Briefly, transverse vibratome sections were subjected to EdU detection using the Click-iT EdU Cell Proliferation Kit (ThermoFisher), then immunostained with SOX2, MYO7A, and DAPI (Scheibinger et al., 2021b). The sections were cleared and then imaged at 1.0x zoom with a confocal microscope at 40X magnification (Zeiss LSM880; Plan-Apochromat 1.3 numerical aperture, oil immersion) using Zen Black acquisition software at a voxel size of 0.208 × 0.208 × 0.371 μm, and z-depth of 80 μm. The image data were imported into syGlass software (Pidhorskyi et al., 2018) which interfaces with SteamVR tracking and the Oculus Rift virtual reality headset, touch controllers, and constellation sensors. Using the “count” function, individual cells were manually annotated in 3D virtual reality. The x, y, z coordinates of each count was exported into Vaa3D (**Fig. 4B**), GraphPad Prism v9 (for bar chart in **Fig. 4C**), and MATLAB vR2017b (for 3D graphing using the scatter3 function – **Fig. 4D**).

### Epithelial Cell Cultures and Quantitative RT-PCR

Culturing of sensory epithelia was conducted as previously described for the utricle (Burns et al., 2008), with some adaptations. 14 mm MatTek dishes were coated with CellTak according to the manufacturer’s instructions, allowing 20 min for absorption, and washing afterwards with sterile water. Peeled sensory epithelia in Medium 199 + 10 μg/mL ciprofloxacin (no serum) were mouth pipetted onto the MatTek dish. The microwell was then filled with the same culture medium. The basilar papilla epithelia were arranged with an eyebrow such that the hair bundles were facing up. A circular 12 mm coverglass was placed over the microwell of each dish. The dish was centrifuged at 400 g for 10 minutes in an Eppendorf 5810R centrifuge to promote adhesion of the sensory epithelia. After centrifugation, the entire dish was slowly filled with Medium 199 + cipro (no serum), the circular coverglass was carefully removed, and the sensory epithelia were incubated for 1 hour at 39°C in 5% CO_2_. This media was then replaced with sisomicin (0.1 mg/mL), sisomicin + JAK/STAT inhibitor (500 nM), or control vehicle in Medium 199 + cipro + 10% FBS, and incubated overnight at 39°C in 5% CO_2_. The following day, the media was replaced without sisomin, but with the continuation of inhibitors for another day.

Total RNA was extracted using the RNAqueous-Micro Kit (ThermoFisher) and reverse transcribed into cDNA using SuperScript IV (Invitrogen). QPCR was performed using Bio-Rad CFX96 Touch Real-Time PCR machine with primer sets listed in **Table S13** and SYBR green detection. Each primer set amplified a single band as determined by gel electrophoresis and melting curve analysis. QPCR data were analyzed using the 2^−ΔΔCt^ method (Livak and Schmittgen, 2001) relative to HSPA8 (housekeeping gene). Statistics were conducted in GraphPad Prism version 9.1.2.

## References

Alizadeh, A., Dyck, S. M. and Karimi-Abdolrezaee, S. (2019). Traumatic Spinal Cord Injury: An Overview of Pathophysiology, Models and Acute Injury Mechanisms. Front. Neurol. 10, 282.

Aruga, J. and Mikoshiba, K. (2003). Identification and characterization of Slitrk, a novel neuronal transmembrane protein family controlling neurite outgrowth. Mol. Cell. Neurosci. 24, 117–129.

Bacher, R., Chu, L.-F., Leng, N., Gasch, A. P., Thomson, J. A., Stewart, R. M., Newton, M. and Kendziorski, C. (2017). SCnorm: robust normalization of single-cell RNA-seq data. Nat. Methods 14, 584–586.

Basters, A., Knobeloch, K. and Fritz, G. (2018). How USP18 deals with ISG15-modified proteins: structural basis for the specificity of the protease. FEBS J. 285, 1024–1029.

Benkafadar, N., Janesick, A., Scheibinger, M., Ling, A. H., Jan, T. A. and Heller, S. (2021). Transcriptomic characterization of dying hair cells in the avian cochlea. Cell Rep. 34, 108902.

Bermingham, N. A. (1999). Math1: An Essential Gene for the Generation of Inner Ear Hair Cells. Science 284, 1837–1841.

Bermingham, N. A., Hassan, B. A., Price, S. D., Vollrath, M. A., Ben-Arie, N., Eatock, R. A., Bellen, H. J., Lysakowski, A. and Zoghbi, H. Y. (1999). Math1: An Essential Gene for the Generation of Inner Ear Hair Cells. Science 284, 1837.

Bermingham-McDonogh, O. and Rubel, E. W. (2003). Hair cell regeneration: winging our way towards a sound future. Curr. Opin. Neurobiol. 13, 119–126.

Bhave, S. A., Oesterle, E. C. and Coltrera, M. D. (1998). Macrophage and microglia-like cells in the avian inner ear. J. Comp. Neurol. 398, 241–256.

Brockes, J. P. and Kumar, A. (2008). Comparative Aspects of Animal Regeneration. Annu. Rev. Cell Dev. Biol. 24, 525–549.

Brose, K. and Tessier-Lavigne, M. (2000). Slit proteins: key regulators of axon guidance, axonal branching, and cell migration. Curr. Opin. Neurobiol. 10, 95–102.

Burns, J., Christophel, J. J., Collado, M. S., Magnus, C., Carfrae, M. and Corwin, J. T. (2008). Reinforcement of cell junctions correlates with the absence of hair cell regeneration in mammals and its occurrence in birds. J. Comp. Neurol. 511, 396–414.

Burns, J. C., Kelly, M. C., Hoa, M., Morell, R. J. and Kelley, M. W. (2015). Single-cell RNA-Seq resolves cellular complexity in sensory organs from the neonatal inner ear. Nat. Commun. 6, 8557.

Cafaro, J., Lee, G. S. and Stone, J. S. (2007). Atoh1 expression defines activated progenitors and differentiating hair cells during avian hair cell regeneration. Dev. Dyn. 236, 156–170.

Calarco, J. A., Superina, S., O’Hanlon, D., Gabut, M., Raj, B., Pan, Q., Skalska, U., Clarke, L., Gelinas, D., van der Kooy, D., et al. (2009). Regulation of Vertebrate Nervous System Alternative Splicing and Development by an SR-Related Protein. Cell 138, 898–910.

Camp, A. J. and Wijesinghe, R. (2009). Calretinin: Modulator of neuronal excitability. Int. J. Biochem. Cell Biol. 41, 2118–2121.

Chen, Y., Lun, A. T. L. and Smyth, G. K. (2016). From reads to genes to pathways: differential expression analysis of RNA-Seq experiments using Rsubread and the edgeR quasi-likelihood pipeline. F1000Research 5, 1438.

Chen, S., Zhou, Y., Chen, Y. and Gu, J. (2018). fastp: an ultra-fast all-in-one FASTQ preprocessor. Bioinformatics 34, i884–i890.

Choudhury, N. R., Nowak, J. S., Zuo, J., Rappsilber, J., Spoel, S. H. and Michlewski, G. (2014). Trim25 Is an RNA-Specific Activator of Lin28a/TuT4-Mediated Uridylation. Cell Rep. 9, 1265–1272.

Costa, A. and Henrique, D. (2015). Transcriptome profiling of induced hair cells (iHCs) generated by combined expression of Gfi1, Pou4f3 and Atoh1 during embryonic stem cell differentiation. Genomics Data 6, 77–80.

Dechesne, C. J., Rabejac, D. and Desmadryl, G. (1994). Development of calretinin immunoreactivity in the mouse inner ear. J. Comp. Neurol. 346, 517–529.

Dedoni, S., Olianas, M. C., Ingianni, A. and Onali, P. (2012). Type I interferons impair BDNF-induced cell signaling and neurotrophic activity in differentiated human SH-SY5Y neuroblastoma cells and mouse primary cortical neurons: Type I interferon inhibition of BDNF signaling. J. Neurochem. 122, 58–71.

Dijkmans, T., Antonia van Hooijdonk, L., Fitzsimons, C. and Vreugdenhil, E. (2010). The Doublecortin Gene Family and Disorders of Neuronal Structure. Cent. Nerv. Syst. Agents Med. Chem. 10, 32–46.

Ding, E., Liu, J., Guo, H., Shen, H., Zhang, H., Gong, W., Song, H. and Zhu, B. (2018). DNMT1 and DNMT3A haplotypes associated with noise-induced hearing loss in Chinese workers. Sci. Rep. 8, 12193.

Dinh, C. T., Goncalves, S., Bas, E., Van De Water, T. R. and Zine, A. (2015). Molecular regulation of auditory hair cell death and approaches to protect sensory receptor cells and/or stimulate repair following acoustic trauma. Front. Cell. Neurosci. 9,.

Dobin, A., Davis, C. A., Schlesinger, F., Drenkow, J., Zaleski, C., Jha, S., Batut, P., Chaisson, M. and Gingeras, T. R. (2013). STAR: ultrafast universal RNA-seq aligner. Bioinformatics 29, 15–21.

Dooling, R. J., Dent, M. L., Lauer, A. M. and Ryals, B. M. (2008). Functional Recovery After Hair Cell Regeneration in Birds. In Hair Cell Regeneration, Repair, and Protection (ed. Salvi, R. J.), Popper, A. N.), and Fay, R. R.), pp. 117–140. New York, NY: Springer New York.

Du, T.-T., Dewey, J. B., Wagner, E. L., Cui, R., Heo, J., Park, J.-J., Francis, S. P., Perez-Reyes, E., Guillot, S. J., Sherman, N. E., et al. (2019). LMO7 deficiency reveals the significance of the cuticular plate for hearing function. Nat. Commun. 10, 1117.

Duckert, L. G. and Rubel, E. W. (1990). Ultrastructural observations on regenerating hair cells in the chick basilar papilla. Hear. Res. 48, 161–182.

Duckert, L. G. and Rubel, E. W. (1993). Morphological correlates of functional recovery in the chicken inner ear after gentamycin treatment. J. Comp. Neurol. 331, 75–96.

Eddleston, M. and Mucke, L. (1993). Molecular profile of reactive astrocytes—Implications for their role in neurologic disease. Neuroscience 54, 15–36.

Edmonds, B., Reyes, R., Schwaller, B. and Roberts, W. M. (2000). Calretinin modifies presynaptic calcium signaling in frog saccular hair cells. Nat. Neurosci. 3, 786–790.

Ellwanger, D. C., Scheibinger, M., Dumont, R. A., Barr-Gillespie, P. G. and Heller, S. (2018). Transcriptional Dynamics of Hair-Bundle Morphogenesis Revealed with CellTrails. Cell Rep. 23, 2901–2914.e13.

Fettiplace, R. and Kim, K. X. (2014). The Physiology of Mechanoelectrical Transduction Channels in Hearing. Physiol. Rev. 94, 951–986.

Fischer, F. P. (1992). Quantitative analysis of the innervation of the chicken basilar papilla. Hear. Res. 61, 167–178.

Franciosi, S., Choi, H. B., Kim, S. U. and McLarnon, J. G. (2002). Interferon-gamma acutely induces calcium influx in human microglia. J. Neurosci. Res. 69, 607–613.

Francis, S. P. and Cunningham, L. L. (2017). Non-autonomous Cellular Responses to Ototoxic Drug-Induced Stress and Death. Front. Cell. Neurosci. 11, 252.

Fujitani, M., Yamagishi, S., Che, Y. H., Hata, K., Kubo, T., Ino, H., Tohyama, M. and Yamashita, T. (2004). P311 accelerates nerve regeneration of the axotomized facial nerve. J. Neurochem. 91, 737–744.

Gall, D., Roussel, C., Susa, I., D’Angelo, E., Rossi, P., Bearzatto, B., Galas, M. C., Blum, D., Schurmans, S. and Schiffmann, S. N. (2003). Altered Neuronal Excitability in Cerebellar Granule Cells of Mice Lacking Calretinin. J. Neurosci. 23, 9320–9327.

Gillespie, P. G. and Müller, U. (2009). Mechanotransduction by Hair Cells: Models, Molecules, and Mechanisms. Cell 139, 33–44.

Girod, D. A., Tucci, D. L. and Rubel, E. W. (1991). Anatomical Correlates of Functional Recovery in the Avian Inner Ear Following Aminoglycoside Ototoxicity: The Laryngoscope 101, 1139–1149.

Goodyear, R. and Richardson, G. (1997). Pattern Formation in the Basilar Papilla: Evidence for Cell Rearrangement. J. Neurosci. 17, 6289–6301.

Goossens, K. E., Ward, A. C., Lowenthal, J. W. and Bean, A. G. D. (2013). Chicken interferons, their receptors and interferon-stimulated genes. Dev. Comp. Immunol. 41, 370–376.

Gordon, C. T., Wade, C., Brinas, I. and Farlie, P. G. (2011). CXCL14 expression during chick embryonic development. Int. J. Dev. Biol. 55, 335–340.

Gubbels, S. P., Woessner, D. W., Mitchell, J. C., Ricci, A. J. and Brigande, J. V. (2008). Functional auditory hair cells produced in the mammalian cochlea by in utero gene transfer. Nature 455, 537–541.

Hackney, C. M., Mahendrasingam, S., Jones, E. M. C. and Fettiplace, R. (2003). The Distribution of Calcium Buffering Proteins in the Turtle Cochlea. J. Neurosci. 23, 4577–4589.

Harper, J. L., Wilson, T. E. and Mitchell, R. M. (2020). Case report of two children with auditory neuropathy spectrum disorder related to a neurofascin (NFASC) gene variant. Int. J. Pediatr. Otorhinolaryngol. 131, 109863.

Hatten, M. E., Liem, R. K. H., Shelanski, M. L. and Mason, C. A. (1991). Astroglia in CNS injury. Glia 4, 233–243.

Hawkins, R. D., Helms, C. A., Winston, J. B., Warchol, M. E. and Lovett, M. (2006). Applying genomics to the avian inner ear: Development of subtractive cDNA resources for exploring sensory function and hair cell regeneration. Genomics 87, 801–808.

Hayashi, Y., Suzuki, H., Nakajima, W., Uehara, I., Tanimura, A., Himeda, T., Koike, S., Katsuno, T., Kitajiri, S., Koyanagi, N., et al. (2020). Cochlear supporting cells function as macrophage-like cells and protect audiosensory receptor hair cells from pathogens. Sci. Rep. 10, 6740.

Heller, S., Bell, A. M., Denis, C. S., Choe, Y. and Hudspeth, A. J. (2002). Parvalbumin 3 is an Abundant Ca2+ Buffer in Hair Cells. JARO - J. Assoc. Res. Otolaryngol. 3, 488–498.

Hirose, K., Rutherford, M. A. and Warchol, M. E. (2017). Two cell populations participate in clearance of damaged hair cells from the sensory epithelia of the inner ear. Hear. Res. 352, 70–81.

Hou, J., Han, L., Zhao, Z., Liu, H., Zhang, L., Ma, C., Yi, F., Liu, B., Zheng, Y. and Gao, C. (2021). USP18 positively regulates innate antiviral immunity by promoting K63-linked polyubiquitination of MAVS. Nat. Commun. 12, 2970.

Hu, Z. and Corwin, J. T. (2007). Inner ear hair cells produced in vitro by a mesenchymal-to-epithelial transition. Proc. Natl. Acad. Sci. 104, 16675–16680.

Hu, B. H., Zhang, C. and Frye, M. D. (2018). Immune cells and non-immune cells with immune function in mammalian cochleae. Hear. Res. 362, 14–24.

Jahan, I., Pan, N., Kersigo, J. and Fritzsch, B. (2010). Neurod1 Suppresses Hair Cell Differentiation in Ear Ganglia and Regulates Hair Cell Subtype Development in the Cochlea. PLoS ONE 5, e11661.

Janesick, A. S. and Heller, S. (2019). Stem Cells and the Bird Cochlea—Where Is Everybody? Cold Spring Harb. Perspect. Med. 9, a033183.

Janesick, A., Scheibinger, M. and Heller, S. (2021a). In vivo hair cell damage model and new molecular tools to study regeneration of the avian cochlea. In Developmental, Physiological and Functional Neurobiology of the Inner Ear, p. Springer (In Press).

Janesick, A., Scheibinger, M., Benkafadar, N., Kirti, S., Ellwanger, D. C. and Heller, S. (2021b). Cell-type identity of the avian cochlea. Cell Rep. 34, 108900.

Jassal, B., Matthews, L., Viteri, G., Gong, C., Lorente, P., Fabregat, A., Sidiropoulos, K., Cook, J., Gillespie, M., Haw, R., et al. (2019). The reactome pathway knowledgebase. Nucleic Acids Res. gkz1031.

Jiang, H., Patel, P. H., Kohlmaier, A., Grenley, M. O., McEwen, D. G. and Edgar, B. A. (2009). Cytokine/Jak/Stat Signaling Mediates Regeneration and Homeostasis in the Drosophila Midgut. Cell 137, 1343–1355.

Julier, Z., Park, A. J., Briquez, P. S. and Martino, M. M. (2017). Promoting tissue regeneration by modulating the immune system. Acta Biomater. 53, 13–28.

Katayama, K., Zine, A., Ota, M., Matsumoto, Y., Inoue, T., Fritzsch, B. and Aruga, J. (2009). Disorganized Innervation and Neuronal Loss in the Inner Ear of Slitrk6-Deficient Mice. PLoS ONE 4, e7786.

Kaur, T., Zamani, D., Tong, L., Rubel, E. W., Ohlemiller, K. K., Hirose, K. and Warchol, M. E. (2015). Fractalkine Signaling Regulates Macrophage Recruitment into the Cochlea and Promotes the Survival of Spiral Ganglion Neurons after Selective Hair Cell Lesion. J. Neurosci. 35, 15050–15061.

Klein, C. J., Botuyan, M.-V., Wu, Y., Ward, C. J., Nicholson, G. A., Hammans, S., Hojo, K., Yamanishi, H., Karpf, A. R., Wallace, D. C., et al. (2011). Mutations in DNMT1 cause hereditary sensory neuropathy with dementia and hearing loss. Nat. Genet. 43, 595–600.

Kolla, L., Kelly, M. C., Mann, Z. F., Anaya-Rocha, A., Ellis, K., Lemons, A., Palermo, A. T., So, K. S., Mays, J. C., Orvis, J., et al. (2020). Characterization of the development of the mouse cochlear epithelium at the single cell level. Nat. Commun. 11, 2389.

Krüger, M., Schmid, T., Krüger, S., Bober, E. and Braun, T. (2006). Functional redundancy of NSCL-1 and NeuroD during development of the petrosal and vestibulocochlear ganglia. Eur. J. Neurosci. 24, 1581–1590.

Ladrech, S., Wang, J., Simonneau, L., Puel, J.-L. and Lenoir, M. (2007). Macrophage contribution to the response of the rat organ of Corti to amikacin. J. Neurosci. Res. 85, 1970–1979.

Ladrech, S., Eybalin, M., Puel, J.-L. and Lenoir, M. (2017). Epithelial–mesenchymal transition, and collective and individual cell migration regulate epithelial changes in the amikacin-damaged organ of Corti. Histochem. Cell Biol. 148, 129–142.

Lee, S., Kurioka, T., Lee, M. Y., Beyer, L. A., Swiderski, D. L., Ritter, K. E. and Raphael, Y. (2021). Scar Formation and Debris Elimination during Hair Cell Degeneration in the Adult DTR Mouse. Neuroscience 453, 57–68.

Li, X.-J. and Doetzlhofer, A. (2020). LIN28B/let-7 control the ability of neonatal murine auditory supporting cells to generate hair cells through mTOR signaling. Proc. Natl. Acad. Sci. 117, 22225–22236.

Liang, J., Wang, D., Renaud, G., Wolfsberg, T. G., Wilson, A. F. and Burgess, S. M. (2012). The stat3/socs3a Pathway Is a Key Regulator of Hair Cell Regeneration in Zebrafish stat3/socs3a Pathway: Regulator of Hair Cell Regeneration. J. Neurosci. 32, 10662–10673.

Lin, X., Zhou, Q., Zhao, C., Lin, G., Xu, J. and Wen, Z. (2019). An Ectoderm-Derived Myeloid-like Cell Population Functions as Antigen Transporters for Langerhans Cells in Zebrafish Epidermis. Dev. Cell 49, 605–617.e5.

Lipkowitz, S., Göbel, V., Varterasian, M. L., Nakahara, K., Tchorz, K. and Kirsch, I. R. (1992). A comparative structural characterization of the human NSCL-1 and NSCL-2 genes. Two basic helix-loop-helix genes expressed in the developing nervous system. J. Biol. Chem. 267, 21065–21071.

Liu, H., Gao, W., Wen, W. and Zhang, Y. (2006). Taurine modulates calcium influx through L-type voltage-gated calcium channels in isolated cochlear outer hair cells in guinea pigs. Neurosci. Lett. 399, 23–26.

Liu, P., Ge, X., Ding, H., Jiang, H., Christensen, B. M. and Li, J. (2012). Role of Glutamate Decarboxylase-like Protein 1 (GADL1) in Taurine Biosynthesis. J. Biol. Chem. 287, 40898–40906.

Livak, K. J. and Schmittgen, T. D. (2001). Analysis of Relative Gene Expression Data Using Real-Time Quantitative PCR and the 2−ΔΔCT Method. Methods 25, 402–408.

Lorenzen, S. M., Duggan, A., Osipovich, A. B., Magnuson, M. A. and García-Añoveros, J. (2015). Insm1 promotes neurogenic proliferation in delaminated otic progenitors. Mech. Dev. 138, 233–245.

Mahootchi, E., Cannon Homaei, S., Kleppe, R., Winge, I., Hegvik, T.-A., Megias-Perez, R., Totland, C., Mogavero, F., Baumann, A., Glennon, J. C., et al. (2020). GADL1 is a multifunctional decarboxylase with tissue-specific roles in β-alanine and carnosine production. Sci. Adv. 6, eabb3713.

Malakhova, O. A., Kim, K. I. I., Luo, J.-K., Zou, W., Kumar, K. G. S., Fuchs, S. Y., Shuai, K. and Zhang, D.-E. (2006). UBP43 is a novel regulator of interferon signaling independent of its ISG15 isopeptidase activity. EMBO J. 25, 2358–2367.

Martín-Vicente, M., Medrano, L. M., Resino, S., García-Sastre, A. and Martínez, I. (2017). TRIM25 in the Regulation of the Antiviral Innate Immunity. Front. Immunol. 8, 1187.

Matsunaga, M., Kita, T., Yamamoto, R., Yamamoto, N., Okano, T., Omori, K., Sakamoto, S. and Nakagawa, T. (2020). Initiation of Supporting Cell Activation for Hair Cell Regeneration in the Avian Auditory Epithelium: An Explant Culture Model. Front. Cell. Neurosci. 14, 583994.

McElrath, C., Espinosa, V., Lin, J.-D., Peng, J., Sridhar, R., Dutta, O., Tseng, H.-C., Smirnov, S. V., Risman, H., Sandoval, M. J., et al. (2021). Critical role of interferons in gastrointestinal injury repair. Nat. Commun. 12, 2624.

McIntyre, J. C., Titlow, W. B. and McClintock, T. S. (2010). Axon growth and guidance genes identify nascent, immature, and mature olfactory sensory neurons: Developmental Regulation of Axon Guidance Genes. J. Neurosci. Res. 88, 3243–3256.

Menendez, L., Trecek, T., Gopalakrishnan, S., Tao, L., Markowitz, A. L., Yu, H. V., Wang, X., Llamas, J., Huang, C., Lee, J., et al. (2020). Generation of inner ear hair cells by direct lineage conversion of primary somatic cells. eLife 9, e55249.

Moisan, A., Villa-Vialaneix, N. and Gonzales, I. (2014). Practical statistical analysis of RNA-Seq data - edgeR.

Monzack, E. L. and Cunningham, L. L. (2013). Lead roles for supporting actors: Critical functions of inner ear supporting cells. Hear. Res. 303, 20–29.

Munnamalai, V., Sienknecht, U. J., Duncan, R. K., Scott, M. K., Thawani, A., Fantetti, K. N., Atallah, N. M., Biesemeier, D. J., Song, K. H., Luethy, K., et al. (2017). Wnt9a Can Influence Cell Fates and Neural Connectivity across the Radial Axis of the Developing Cochlea. J. Neurosci. 37, 8975–8988.

Murdoch, J., Eddleston, J., Leblond-Bourget, N., Stanier, P. and Copp, A. (1999). Sequence and expression analysis of Nhlh1: a basic helix-loop-helix gene implicated in neurogenesis. Dev Genet 24, 165–177.

Nagao, M., Ogata, T., Sawada, Y. and Gotoh, Y. (2016). Zbtb20 promotes astrocytogenesis during neocortical development. Nat. Commun. 7, 11102.

Neerukonda, S. N. and Katneni, U. (2020). Avian Pattern Recognition Receptor Sensing and Signaling. Vet. Sci. 7, 14.

Nguyen, A. V. and Soulika, A. M. (2019). The Dynamics of the Skin’s Immune System. Int. J. Mol. Sci. 20, 1811.

O’Halloran, E. K. and Oesterle, E. C. (2004). Characterization of leukocyte subtypes in chicken inner ear sensory epithelia. J. Comp. Neurol. 475, 340–360.

Ordonez, J. L. and Tekin, M. (2017). Deafness and Myopia Syndrome. University of Washington, Seattle, Seattle (WA).

Orvis, J., Gottfried, B., Kancherla, J., Adkins, R. S., Song, Y., Dror, A. A., Olley, D., Rose, K., Chrysostomou, E., Kelly, M. C., et al. (2021). gEAR: Gene Expression Analysis Resource portal for community-driven, multi-omic data exploration. Nat. Methods.

Oshima, K., Shin, K., Diensthuber, M., Peng, A. W., Ricci, A. J. and Heller, S. (2010). Mechanosensitive Hair Cell-like Cells from Embryonic and Induced Pluripotent Stem Cells. Cell 141, 704–716.

Picelli, S., Faridani, O. R., Björklund, Å. K., Winberg, G., Sagasser, S. and Sandberg, R. (2014). Full-length RNA-seq from single cells using Smart-seq2. Nat. Protoc. 9, 171–181.

Pidhorskyi, S., Morehead, M., Jones, Q., Spirou, G. and Doretto, G. (2018). syGlass: Interactive Exploration of Multidimensional Images Using Virtual Reality Head-mounted Displays. ArXiv180408197 Cs.

Pilipenko, V. V., Reece, A., Choo, D. I. and Greinwald, J. H. (2004). Genomic organization and expression analysis of the murine Fam3c gene. Gene 335, 159–168.

Quesnel-Vallières, M., Irimia, M., Cordes, S. P. and Blencowe, B. J. (2015). Essential roles for the splicing regulator nSR100/SRRM4 during nervous system development. Genes Dev. 29, 746–759.

Quintás-Cardama, A., Vaddi, K., Liu, P., Manshouri, T., Li, J., Scherle, P. A., Caulder, E., Wen, X., Li, Y., Waeltz, P., et al. (2010). Preclinical characterization of the selective JAK1/2 inhibitor INCB018424: therapeutic implications for the treatment of myeloproliferative neoplasms. Blood 115, 3109–3117.

Rai, V., Wood, M. B., Feng, H., Schabla, Nathan. M., Tu, S. and Zuo, J. (2020). The immune response after noise damage in the cochlea is characterized by a heterogeneous mix of adaptive and innate immune cells. Sci. Rep. 10, 15167.

Raphael, Y., Kim, Y.-H., Osumi, Y. and Izumikawa, M. (2007). Non-sensory cells in the deafened organ of Corti: approaches for repair. Int. J. Dev. Biol. 51, 649–654.

Ratié, L., Ware, M., Barloy-Hubler, F., Romé, H., Gicquel, I., Dubourg, C., David, V. and Dupé, V. (2013). Novel genes upregulated when NOTCH signalling is disrupted during hypothalamic development. Neural Develop. 8, 25.

Ratié, L., Ware, M., Jagline, H., David, V. and Dupé, V. (2014). Dynamic expression of Notch-dependent neurogenic markers in the chick embryonic nervous system. Front. Neuroanat. 8,.

Rau, A., Gallopin, M., Celeux, G. and Jaffrézic, F. (2013). Data-based filtering for replicated high-throughput transcriptome sequencing experiments. Bioinformatics 29, 2146–2152.

Rio, C., Dikkes, P., Liberman, M. C. and Corfas, G. (2002). Glial fibrillary acidic protein expression and promoter activity in the inner ear of developing and adult mice. J. Comp. Neurol. 442, 156–162.

Robinson, M. D. and Oshlack, A. (2010). A scaling normalization method for differential expression analysis of RNA-seq data. Genome Biol. 11, R25.

Robinson, M. D., McCarthy, D. J. and Smyth, G. K. (2010). edgeR: a Bioconductor package for differential expression analysis of digital gene expression data. Bioinformatics 26, 139–140.

Ryals, B. M., Dent, M. L. and Dooling, R. J. (2013). Return of function after hair cell regeneration. Hear. Res. 297, 113–120.

Sánchez-Solana, B., Nueda, M. L., Ruvira, M. D., Ruiz-Hidalgo, M. J., Monsalve, E. M., Rivero, S., García-Ramírez, J. J., Díaz-Guerra, M. J. M., Baladrón, V. and Laborda, J. (2011). The EGF-like proteins DLK1 and DLK2 function as inhibitory non-canonical ligands of NOTCH1 receptor that modulate each other’s activities. Biochim. Biophys. Acta BBA - Mol. Cell Res. 1813, 1153–1164.

Santhakumar, D., Rubbenstroth, D., Martinez-Sobrido, L. and Munir, M. (2017). Avian Interferons and Their Antiviral Effectors. Front. Immunol. 8,.

Saunders, J. C. and Salvi, R. J. (2008). Recovery of Function in the Avian Auditory System After Ototrauma. In Hair Cell Regeneration, Repair, and Protection (ed. Salvi, R. J.), Popper, A. N.), and Fay, R. R.), pp. 77–116. New York, NY: Springer New York.

Sayyid, Z. N., Wang, T., Chen, L., Jones, S. M. and Cheng, A. G. (2019). Atoh1 Directs Regeneration and Functional Recovery of the Mature Mouse Vestibular System. Cell Rep. 28, 312–324.e4.

Scheffer, D., Sage, C., Plazas, P. V., Huang, M., Wedemeyer, C., Zhang, D.-S., Chen, Z.-Y., Elgoyhen, A. B., Corey, D. P. and Pingault, V. (2007). The α1 subunit of nicotinic acetylcholine receptors in the inner ear: transcriptional regulation by ATOH1 and co-expression with the γ subunit in hair cells. J. Neurochem. 0, 071027034430002-???

Scheibinger, M., Ellwanger, D. C., Corrales, C. E., Stone, J. S. and Heller, S. (2018). Aminoglycoside Damage and Hair Cell Regeneration in the Chicken Utricle. J. Assoc. Res. Otolaryngol. 19, 17–29.

Scheibinger, M., Janesick, A., Benkafadar, N., Ellwanger, D. C., Jan, T. A. and Heller, S. (2021a). Cell-Type Identity of the Avian Utricle. SSRN Electron. J.

Scheibinger, M., Janesick, A., Diaz, G. and Heller, S. (2021b). Immunohistochemistry and In Situ mRNA Detection using Inner Ear Vibratome Sections. In Developmental, Physiological and Functional Neurobiology of the Inner Ear, p. Springer (In Press).

Schwaller, B. (2009). The continuing disappearance of “pure” Ca2+ buffers. Cell. Mol. Life Sci. 66, 275–300.

Schwaller, B. and Herrmann, B. (1997). Regulated redistribution of calretinins in WiDr cells. Cell Death Differ. 4, 325–333.

Shrestha, B. R., Chia, C., Wu, L., Kujawa, S. G., Liberman, M. C. and Goodrich, L. V. (2018). Sensory Neuron Diversity in the Inner Ear Is Shaped by Activity. Cell 174, 1229–1246.e17.

Stone, J. S. and Rubel, E. W. (1999). Delta1 expression during avian hair cell regeneration. Development 126, 961–973.

Stone, J. S. and Rubel, E. W. (2000). Cellular studies of auditory hair cell regeneration in birds. Proc. Natl. Acad. Sci. 97, 11714–11721.

Su, S., Law, C. W., Ah-Cann, C., Asselin-Labat, M.-L., Blewitt, M. E. and Ritchie, M. E. (2017). Glimma: interactive graphics for gene expression analysis. Bioinformatics 33, 2050–2052.

Sun, L., Miyoshi, H., Origanti, S., Nice, T. J., Barger, A. C., Manieri, N. A., Fogel, L. A., French, A. R., Piwnica-Worms, D., Piwnica-Worms, H., et al. (2015). Type I Interferons Link Viral Infection to Enhanced Epithelial Turnover and Repair. Cell Host Microbe 17, 85–97.

Sun, S., Babola, T., Pregernig, G., So, K. S., Nguyen, M., Su, S.-S. M., Palermo, A. T., Bergles, D. E., Burns, J. C. and Müller, U. (2018). Hair Cell Mechanotransduction Regulates Spontaneous Activity and Spiral Ganglion Subtype Specification in the Auditory System. Cell 174, 1247–1263.e15.

Talman, V. and Ruskoaho, H. (2016). Cardiac fibrosis in myocardial infarction—from repair and remodeling to regeneration. Cell Tissue Res. 365, 563–581.

Tornabene, S. V., Sato, K., Pham, L., Billings, P. and Keithley, E. M. (2006). Immune cell recruitment following acoustic trauma. Hear. Res. 222, 115–124.

Tran, A. P., Warren, P. M. and Silver, J. (2018). The Biology of Regeneration Failure and Success After Spinal Cord Injury. Physiol. Rev. 98, 881–917.

Tucci, D. L. and Rubel, E. W. (1990). Physiologic Status of Regenerated Hair Cells in the Avian Inner Ear following Aminoglycoside Ototoxicity. Otolaryngol. Neck Surg. 103, 443–450.

Ulgen, E., Ozisik, O. and Sezerman, O. U. (2019). pathfindR: An R Package for Comprehensive Identification of Enriched Pathways in Omics Data Through Active Subnetworks. Front. Genet. 10, 858.

Wan, G., Corfas, G. and Stone, J. S. (2013). Inner ear supporting cells: Rethinking the silent majority. Semin. Cell Dev. Biol. 24, 448–459.

Warchol, M. (1997). Macrophage activity in organ cultures of the avian cochlea: demonstration of a resident population and recruitment to sites of hair cell lesions. J Neurobiol 33, 724–734.

Warchol, M. E., Schwendener, R. A. and Hirose, K. (2012). Depletion of Resident Macrophages Does Not Alter Sensory Regeneration in the Avian Cochlea. PLoS ONE 7, e51574.

Warchol, M. E., Schrader, A. and Sheets, L. (2021). Macrophages Respond Rapidly to Ototoxic Injury of Lateral Line Hair Cells but Are Not Required for Hair Cell Regeneration. Front. Cell. Neurosci. 14, 613246.

Wilmes, S., Beutel, O., Li, Z., Francois-Newton, V., Richter, C. P., Janning, D., Kroll, C., Hanhart, P., Hötte, K., You, C., et al. (2015). Receptor dimerization dynamics as a regulatory valve for plasticity of type I interferon signaling. J. Cell Biol. 209, 579–593.

Wiwatpanit, T., Lorenzen, S. M., Cantú, J. A., Foo, C. Z., Hogan, A. K., Márquez, F., Clancy, J. C., Schipma, M. J., Cheatham, M. A., Duggan, A., et al. (2018). Trans-differentiation of outer hair cells into inner hair cells in the absence of INSM1. Nature 563, 691–695.

Wood, M. B. and Zuo, J. (2017). The Contribution of Immune Infiltrates to Ototoxicity and Cochlear Hair Cell Loss. Front. Cell. Neurosci. 11,.

Zheng, J. L. and Gao, W.-Q. (1997). Analysis of Rat Vestibular Hair Cell Development and Regeneration Using Calretinin as an Early Marker. J. Neurosci. 17, 8270–8282.

